# Variation in the molecular clock of primates

**DOI:** 10.1101/036434

**Authors:** Priya Moorjani, Carlos Eduardo G. Amorim, Peter F. Arndt, Molly Przeworski

## Abstract

Events in primate evolution are often dated by assuming a “molecular clock”, i.e., a constant rate of substitution per unit time, but the validity of this assumption remains unclear. Among mammals, it is well known that there exists substantial variation in yearly substitution rates. Such variation is to be expected from differences in life-history traits, suggesting that it should also be found among primates. Motivated by these considerations, we analyze whole genomes from ten primate species, including Old World Monkeys (OWMs), New World Monkeys (NWMs) and apes, focusing on putatively neutral autosomal sites and controlling for possible effects of biased gene conversion and methylation at CpG sites. We find that substitution rates are ˜65% higher in lineages leading from the hominoid-NWM ancestor to NWMs than to apes. Within apes, rates are ˜2% higher in chimpanzees and ˜7% higher in the gorilla than in humans. Substitution types subject to biased gene conversion show no more variation among species than those not subject to it. Not all mutation types behave similarly, however: in particular, transitions at CpG sites exhibit a more clock-like behavior than do other types, presumably due to their non-replicative origin. Thus, not only the total rate, but also the mutational spectrum varies among primates. This finding suggests that events in primate evolution are most reliably dated using CpG transitions. Taking this approach, we estimate that the average time to the most recent common ancestor of human and chimpanzee is 12.1 million years and their split time 7.9 million years.

**Significance statement:** Much of our understanding of the chronology of human evolution relies on the “molecular clock”, i.e., a constant rate of substitutions per unit time. To evaluate the validity of this assumption, we analyze whole genome sequences from ten primate species. We find that there is substantial variation in the molecular clock between apes and monkeys, and rates even differ within hominoids. Importantly, not all mutation types behave similarly: notably, transitions at CpG sites exhibit a more clock-like behavior than other substitutions, presumably due to their non-replicative origin. Thus, the mutation spectra, and not just the overall substitution rates, are changing across primates. This finding further suggests that events in primate evolution are most reliably dated using CpG transitions.

## Introduction

Germline mutations are the ultimate source of genetic differences among individuals and species. They are thought to arise from a combination of errors in DNA replication (e.g., the chance misincorporation of a base pair) or damage that is unrepaired by the time of replication (e.g., the spontaneous deamination of methylated CpG sites) (1). If mutations are neutral (i.e., do not affect fitness), then the rate at which they arise will be equal to the substitution rate (2). A key consequence is that if mutation rates remain constant over time, substitution rates should likewise be constant.

This assumption of constancy of substitution rates plays a fundamental role in evolutionary genetics, by providing a molecular clock by which to date events inferred from genetic data (3). Notably, important events in human evolution for which there is no fossil record (e.g., when humans and chimpanzees split, or when anatomically modern humans first left the African subcontinent) are dated using a mutation rate obtained from contemporary pedigrees or phylogenetic analysis, assuming the per year rate has remained unchanged for millions of years (4, 5).

Yet we know from studies of mammalian phylogenies, as well as of other taxa, that there can be substantial variation in substitution rates per unit time (6-8). In particular, there is the well known “generation time effect” on substitution rates, which suggests that species with shorter generation time (i.e. mean age of reproduction) have higher mutation rates (9). For instance, mice have a generation time on the order of months (˜10-12 months) compared to ˜29 years in humans, and a two- to three-fold higher substitution rate per year (9). More generally, in a survey of 32 mammalian species, the strongest predictor of substitution rate variation was the generation time (6).

A generation time effect has also been suggested in humans as the basis for a possible slowdown in the yearly mutation rate towards the present. This conjecture is motivated by the observation that the yearly mutation rate estimated from substitutions accumulated between humans and chimpanzees over millions of years (˜10^-9^ per base pair per year (10, 11)) is more than two-fold higher than the yearly mutation rate estimated from *de novo* mutation events in trios (˜0.4 x 10^-9^ per base pair per year (12, 13)). One possibility is that the generation time has increased towards the present, and led to a decrease in the yearly mutation rate (14).

Whether the association between generation time and substitution rates is causal remains unclear, however, correlated traits such as metabolic rate (15), body size (16) and sperm competition (17), may also affect substitution rates. For instance, the metabolic rate hypothesis posits that species with higher basal metabolic rates are subject to higher rates of oxidative stress and hence have a higher mutation rate (15). Body mass has been shown to be negatively correlated to substitution rates, such that smaller animals tend to have higher substitution rates (15). Sexual selection on mating systems may also affect substitution rates, more intense sperm competition leads to selection for higher sperm counts, leading to more cell divisions per unit time during spermatogenesis and a higher male mutation rate (17).

That said, an effect of life history traits such as generation time on the yearly mutation rate is expected based on first principles, given our understanding of oogenesis and spermatogenesis (18, 19). In mammals, oogonial divisions are completed by the birth of the future mother, whereas the spermatogonial stem cells continue to divide post-puberty (18). Thus, the total number of replication-driven mutations inherited by a diploid offspring accrues in a piecewise linear manner with parental age, with the number depending on the number of cell divisions in each developmental stage as well as the per cell division mutation rates (1, 19). These considerations indicate that changes in generation time, onset of puberty and rate of spermatogenesis should all influence yearly mutation rates (1, 19).

Importantly then, primates are well known to differ with regard to most of these traits. In addition to huge variation in body size and metabolic rates, generation time varies almost ten fold, with the shortest generation time observed in prosimians (˜3 years in galago and mouse lemurs (20)) and the longest generation time observed in humans (˜29 years (21)). Species also differ in the strength of sperm competition and rates of spermatogenesis: monkeys have a shorter spermatogenetic division and thus consequently produce more sperm per unit time than do apes (22). Thus, even if the per cell division mutation rate remained constant, we should expect differences in yearly mutation rates among species.

While the factors discussed thus far apply to all sites, variation in substitution rates among species also depends on the type of mutation and the genomic context (i.e., flanking sequence) in which it occurs (7). For example, in mammals, CpG transitions show the least amount of variation in substitution rates among species (7). A plausible explanation is the source of mutations: transitions at methylated CpG sites are thought to occur primarily through spontaneous deamination; if they arise at a constant rate and their repair is inefficient relative to the cell cycle length, as is thought to be the case, then their mutation rate should depend largely on absolute time rather than on the number of cell divisions (23-25).

In addition, even substitutions that are neutral in their effects on fitness may vary in their rate of accumulation among lineages because of biased gene conversion (BGC), the bias towards strong (S: G or C) rather than weak (W: A or T) bases that occurs in the repair of double strand breaks (26). This phenomenon leads to the increased probability of fixation of S alleles (and loss of W alleles) in regions of higher recombination, and can therefore change substitution rates relative to mutation rates (26, 27). The strength of BGC is a function of the degree of bias, the local recombination rate and the effective population size of the species (26). The latter varies by three to four fold among primates (28), and the fine-scale recombination landscape is also likely to differ substantially across species (29, 30).

Empirically, the extent to which substitution rates vary among primate lineages remains unclear. Kim et al. (2006) compared two hominoids (human and chimpanzee) and two OWMs (baboon and rhesus macaque); assuming that the average divergence time of the two pairs of species is identical, they reported that substitution rates at transitions at non-CpG sites differ by approximately 31% between hominoids and OWM, whereas rates of CpG transitions are almost identical (31). In turn, Elango et al. (2006) found that the human branch is ˜2% shorter than chimpanzees (considering the rates from the human-chimpanzee ancestor), and shorter than gorilla (considering rates from the human-gorilla ancestor) (32). While these comparisons raise the possibility that substitution rates are evolving across primates, they are based on limited data, make strong assumptions about divergence times, and rely on parsimony-based approaches that may underestimate substitution rates for divergent species, notably at CpG sites (33). We therefore revisit these questions using whole genome sequence alignments of ten primates, allowing for variable substitution rates along different lineages and explicitly modeling the context dependency of CpG substitutions.

## Materials and Methods

**Data sets and filtering**. We used the following datasets for our analysis: (a) A 12-primate whole genome sequence alignment, with mouse as an outgroup, which is part of a 100-way mammalian phylogeny, mapped using Multiz (34) (referred to as the Multiz dataset). (b) A seven primate whole genome alignment, mapped using the Enredo-Pecan-Ortheus (EPO) pipeline (35) (referred to the EPO dataset); and (c) High coverage genomes for a human (of European descent) that we sequenced (Note S1), a chimpanzee (Ind-D from (36)) and a gorilla (Delphi from (37); data kindly provided by Tomas Marques-Bonet, Institut Biologia Evolutiva (Universitat Pompeu Fabra/CSIC)) (referred to as the high coverage hominoid dataset). For the EPO dataset (b), we removed duplications using the mafDuplicateFilter from mafTools package (38). This software identifies any duplicated region in the alignment block and only retains the sequence with the highest similarity to the consensus sequence. In (c), the genomes were mapped to the orangutan reference genome (ponAbe2) (39), which should be equidistant to humans and extant African great apes (assuming no variation in substitution rates), using bwa-mem (40) with default parameters and the multi-threading option (-t). The coverage after mapping was as follows: human = 30.21, chimpanzee = 31.23 and gorilla = 32.75. Single Nucleotide polymorphisms (SNP) were called using samtools mpileup (version: 0.1.18-dev) (41) with the -B option (to reduce the number of false SNPs called due to misalignments). The bam files were converted to fasta format using BCFtools and seqtk (part of samtools) and only sites that had a minimum quality score of 30 were retained for further analysis (-q30). As we need haploid genomes in our inference procedure, for each polymorphic site in the high coverage genomes, we randomly sampled one allele, thereby generating a pseudo-haploid genome for each species. These high coverage and high quality fasta files were used for pairwise comparisons of human-chimpanzee and human-gorilla genomes, using the orangutan reference genome as an outgroup.

For the three datasets, we filtered out missing data, i.e., any base pair that was aligned to a gap or a missing site in at least one of the primate species. To minimize the effects of selection acting on the base pairs considered, we limited our analyses to the non-coding, non-conserved and non-repetitive regions of the genome. Table S1 includes the source of all annotations used. For each primate species, we excluded sites with the following annotations:

a. Conserved elements annotated using phastCons (42) based on the multiple alignments of 46 primates (43). These annotations were downloaded from UCSC browser (track: phastConsElements46wayPrimates).
b. Coding exons based on the NCBI RNA reference sequences collection annotation or equivalent. These annotations were downloaded from UCSC browser (track: RefSeq Genes).
c. Transposable elements. As the levels of methylation are higher for repetitive regions than non-repetitive regions of the genome (44), which could lead to differences in mutation rates, we removed the repetitive regions including interspersed nuclear elements (LINE and SINE), DNA repeat elements and Long Terminal Repeat elements identified using RepeatMasker (45).

In some cases, we also excluded sites within CpG islands (CGI). Transitions at CpG sites are thought to primarily occur due to spontaneous deamination at methylated cytosines. However, within CGI, most CpGs are hypomethylated (46). As an illustration, comparison of sperm methylation profiles in humans from (47) showed that only 7.5% of CpG sites in annotated CGI have a methylation level of greater than or equal to 40% whereas the vast majority (84.6%) of CpG sites outside CGI have similar or greater methylation levels (Supplementary figure S1). To focus on a more homogeneous set of methylated CpGs, we therefore excluded CGI from the analysis, unless otherwise specified. CGI annotations were downloaded from UCSC browser (track: CpG Islands) (48).

**Estimating substitution rates**. We used Phylofit (49) to estimate autosomal substitutions from the three datasets described above. To access the robustness of the estimates from Phylofit, we also used an alternative maximum likelihood based approach from (50) for the high coverage hominoid genomes. Both methods require as input the topology of the phylogenetic tree for the species represented in the analysis, which were subsets of the primates included in the Multiz dataset, the EPO dataset or that of hominoids. Because these methods assume a single tree for all sites (i.e., ignore the possibility of incomplete lineage sorting), for species pairs with known and non-negligible incomplete lineage sorting, such as chimpanzees/gorillas and gibbons/orangutans, we considered only one of the two lineages in a given analysis (51).

Phylofit (49) analysis was performed with the expectation maximization algorithm (option -E) with medium precision for convergence. We used the U2S substitution model (the general unrestricted dinucleotide model with strand symmetry) with overlapping tuples to estimate lineage-specific CpG substitution rates and UNREST (the general unrestricted single nucleotide model) to estimate the non-CpG substitution rates. To ensure that the branch lengths across U2S and UNREST are comparable, we ran UNREST with fixed branch lengths that were estimated using U2S. For both internal and external branches, Phylofit outputs both the overall branch lengths, accounting for recurrent substitutions at a site, and posterior expected counts of the number of positions at which the descendant base differs from the ancestral base (option -Z).

We used the posterior counts to estimate the number of substitutions involving transitions and transversions for the following types of sites: ancestrally A or T sites (referred to as A/T), ancestrally G or C sites (G/C), ancestrally CG dinucleotides (CpG) and ancestrally G or C sites that are not part of a CG dinucleotide (non-CpG G/C). Specifically, for each context, we estimated the lineage-specific divergence from any internal node to the terminal node X such that X_A1-_>_A2_ is the number of positions at which the ancestral allele A1 (at the internal node) is estimated to have substituted to allele A2 (at the terminal node) and divided by the total count of ancestral alleles A1 on the lineage X, assuming single-step mutations from A1 to A2, thereby implicitly making a parsimony assumption. To study the effects of biased gene conversion, we similarly estimated the substitution rates for strong (S; G/C) and weak (W; A/T) mutations in different substitution contexts (CpG or non-CpG).

For the high coverage hominoid analysis (dataset (c)), we ran Phylofit five times with five different seeds (using -r and -D options). We observed that the estimates were quite unstable, and in particular that a subset of the runs had substantially lower likelihoods. We interpreted this result as indicating that the method sometimes returns values for a local peak in the likelihood surface. To circumvent this problem, we therefore took the estimate for the run with the highest likelihood. Additionally, we also used the maximum likelihood based approach from (50). This approach uses a probabilistic model for sequence evolution and assumes that all nucleotide substitutions except those occurring in a CpG context evolve independently. This leads to 6 parameters in a reverse complement symmetric analysis or 12 parameters if the complement strands evolve with different rates. Substitutions at C and G in the CpG context have their own rates, which yields 3 or 6 additional parameters in the reverse complement symmetric setting or non-reverse complement symmetric setting, respectively. Because the identity of left neighbor or the right neighbor influences substitution processes (e.g. CpG -> TpG or CpG -> CpA), the maximum likelihood approach computes the evolution of tri-nucleotides. Unlike Phylofit, the maximum likelihood approach does not assume that the nucleotide substitution process is in stationary state. This method was run with multi-threading and strand-asymmetry option to estimate the rate of 12 context-free substitutions (A->[C/T/G], T->[A/C/G], non-CpG C->[A/T/G] and non-CpG G->[A/T/G]) and six CpG substitutions (two CpG transitions: CG->[CA/TG] or four CpG transversions: CG->[CC/GG/CT/AG]). To obtain estimates of the number of transitions and transversions for different ancestral contexts (A/T, CpG and non-CpG G/C), we estimated a weighted average of the rates across symmetric classes of substitutions using the counts of the nucleotide contents in the orangutan genome for normalization.

**Assessing the significance of branch length differences in pairwise comparisons**. To test if the branch lengths estimated by Phylofit differ between two species (species1 and species2), we used a likelihood ratio test where the null model (Ho) was that the number of substitutions on the branch leading to species_1_ (*n*-_1_) and branch leading to species_2_(*n*_2_) from the common ancestor are equal, so the proportion (p) = *n*_1_/*n* = 0.5, where *n* = % + *n*_2_. In the alternative model (H_A_), the number of substitutions on the branch leading to species1 was not equal to the number of substitutions on the branch to species2, i.e p ≠ 0.5. Thus, the likelihood ratio statistic

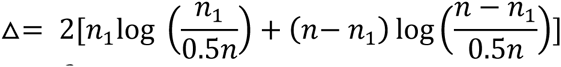

should be approximately *x*^2^ (*df* = 1).

**Estimating the root-leaf variance**. For each substitution type, we constructed a phylogenetic tree using the lineage-specific substitution rates estimated by Phylofit from the Multiz and EPO datasets. We computed the root-leaf distance using the *R* package *adephylo* (52). Following (7), we considered the variance in the root to leaf distance after normalizing by the mean distance.

**Phylogenetically independent contrast analysis:** We tested the correlation between generation time and non-CpG substitution rates using the phylogenetically independent contrasts *(pic)* method described by Felsenstein (1985) (53) implemented in the *R* package *ape* (54). We used the CpG transition rates to define our baseline phylogeny. Generation time estimates assumed for all extant species are shown in Table S2.

**Modeling yearly mutation rates**. To estimate the average yearly mutation rates (μ_y_) for a given set of life-history traits, we used the mutational model from (25). In this model, the mutation rate per year is given by:

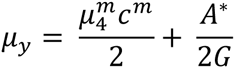

where 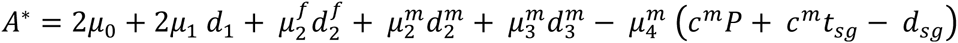. Here, 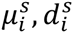 is the mutation rate and number of cell divisions in the *i*^th^ stage of development (*i* = 0: first post-zygotic division, 1: second post-zygotic division to sex differentiation, 3: sex differentiation to birth, 4: birth to puberty and 5: puberty to reproduction) in sex *s* (male or female) respectively, *t_sg_* is the duration of spermatogenesis (in years), *d_sg_* is the number of spermatogonial stem cell divisions required to complete spermatogenesis, 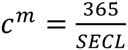 is the number of spermatogonial stem cell divisions each year for a given rate of spermatogenesis (measured by estimating the seminiferous epithelium cycle length (SECL)), *P* is the age of puberty and *G* is the mean age of reproduction (assumed to be the same in males and females).

Following (25), we assumed 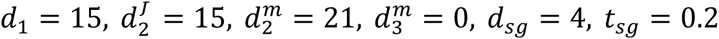 and *μ_2_ = μ_3_* = *μ_4_* = *μ* and *μ_0_ = *μ_1_* = 4μ* to allow for a higher mutation rate in the first two post-zygotic divisions (55). Parameter values for life-history traits used for different species are shown in Table S2.

**Estimating divergence and split times among apes from CpG substitutions**. We estimated the divergence time between human-chimpanzee and human-gorilla using substitutions involving transitions at CpG sites (outside CGI), by:

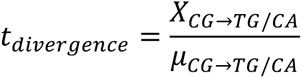

where 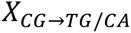 is the expected number of transitions estimated by Phylofit to have occurred at CpG sites on the human lineage since the split from the common ancestor (i.e. either the human-chimpanzee or human-gorilla common ancestor) and 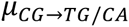 is the estimated per year mutation rate for CpG transitions. The value of 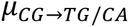 (=3.9x10^-9^ per base pair per year) was obtained by dividing the per generation mutation rate at CpG transitions (= 1.12x10^-7^ per base pair per generation) in (12) by the mean parental age in that study (28.4 years), which is appropriate if the number of CpG transitions is considered linear with age.

As *t_divergence_ = t_*split*_ + t_coalescent_* and assuming a panmictic, constant size population, *t_coalescent_ = 2N_a_ G*,

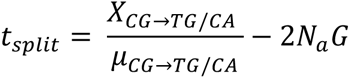

where *N_a_* is effective population size of the ancestral population. We assumed *N_a_* = 5N_h_ (37, 56) and 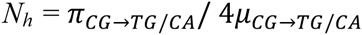, where *N_h_* is the effective population size in humans and 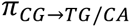 is the average diversity level observed at transitions at CpG sites across 13 contemporary human populations (57, 58).

## Results

We first estimate the number of autosomal substitutions on 10 primate lineages by applying Phylofit (49) to the Multiz sequence alignment (excluding gorilla and gibbon due to concerns about incomplete lineage sorting; see Methods). This method allows us to estimate branch lengths, accounting for uncertainty in the ancestral reconstruction, recurrent substitutions at a site, and allowing for context dependent effects of neighboring nucleotides at CpG dinucleotides (49).

To focus on putatively neutral sites in the genome, in which substitutions more faithfully reflect mutation rates and patterns, we exclude conserved elements, coding exons and transposable elements (referred to as CET in what follows, see Methods). After filtering CET sites and removing missing data, we obtain ˜562 Mb of whole genome sequence alignment across 10 primates. In these data, the total substitution rates vary markedly across the 10 primates (Figure 1). For example, when we compare taxa pairwise, the substitution rates on lineages leading from the hominoid-OWM ancestor to hominoids are on average 2.68% (with a range of 2.63 to 2.74%) whereas rates on lineages leading to OWM are on average 3.66% (3.58 to 3.74%), 1.37-fold higher. These findings are qualitatively consistent with previous, smaller studies (31). Similarly, when considering the distance from the hominoid-NMW ancestor, substitution rates leading to NWM are on average 6.92% (6.89 to 6.94%), 1.64-fold higher than on the lineages leading to hominoids, which are on average 4.23% (4.18 to 4.29%). Substitution rates are also 1.62fold higher in lineages leading to bushbaby (a prosimian) compared to hominoids (Figure 1). However, because of challenges in accurately reconstructing the ancestral state for species that are closer to the outgroup, we believe this estimate to be less reliable and hence do not consider bushbaby in any further analysis.

**Figure 1:**
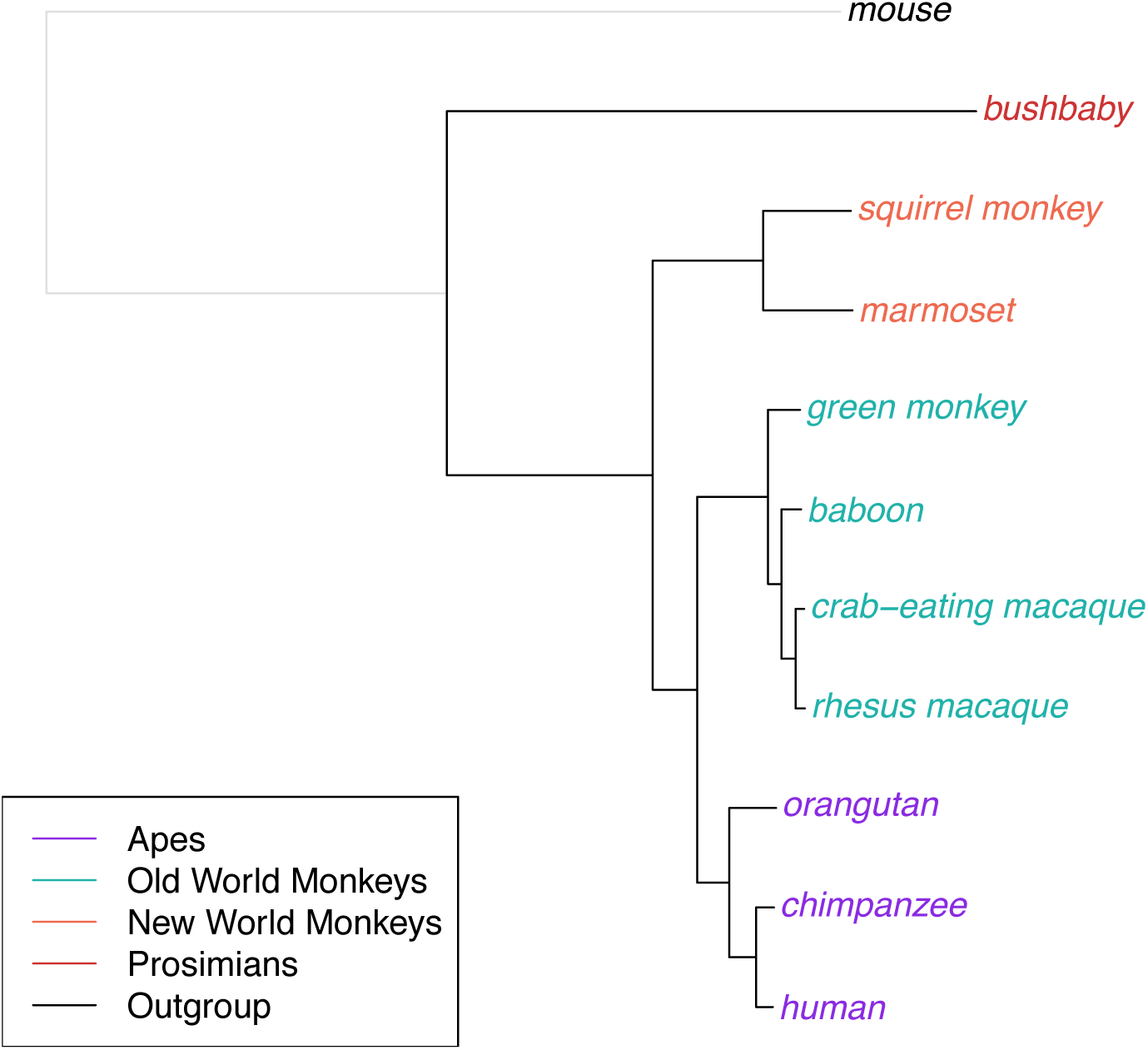
Phylogenetic trees for 10 primates. We estimate neutral substitution rates for 10 primates and an outgroup (mouse, shown in gray) from the Multiz dataset using Phylofit (see Methods for details of data set and filtering). Branch lengths reflect the expected number of neutral substitutions per site along each lineage.

Using bootstrap resampling of 1 Mb regions of the genome suggests tiny standard errors for the substitution rates (e.g., the standard error on lineages leading from the hominoid-OWM ancestor to hominoids is 0.01%), as expected from analyzing such large data sets. These standard errors are likely to be deceptively small, however, as the main source of uncertainty in our analysis is likely due to systematic effects of varying sequence quality, mapping and alignment artifacts among species. To evaluate the impact of these effects, we therefore repeat our analysis using a different sequence alignment of seven primates (the EPO dataset) and apply the same filters. When the species considered are matched between datasets, results are highly similar (Note S2), suggesting results are robust to data quality issues.

To evaluate how substitution patterns differ for mutations generated by distinct mechanisms, we distinguish between transitions at ancestrally CpG sites (referred to as CpG) outside of CpG islands (CGI), which are believed to occur mostly due to the spontaneous deamination of methylated cytosines, and transitions at ancestrally G or C sites outside of a CpG context (referred to as non-CpG G/C), which are thought to primarily occur due to replication errors. (Because CGI are often hypomethylated, we remove these regions from this analysis, and, unless specified otherwise, we refer to transitions at CpG sites outside of CGIs as “CpG transitions”). Mathematical modeling of different substitution mechanisms predicts that mutations that are non-replicative in origin and highly inefficiently repaired should depend on absolute time rather than on the number of cell divisions and hence should be more clock-like among species (25). In contrast, mutations that arise from replication errors or are non-replicative in origin but well repaired should depend on the generation time and other life history traits, and thus their substitution rates could vary considerably across primates (25, 59). Thus, a priori, we expect CpG transitions outside CGI to be more clock-like than other types of substitutions (assuming similar rates of deamination).

In order for our comparisons to not be confounded by biased gene conversion, we focus on the comparison of transitions at CpG sites with those occurring from non-CpG G/C sites. Because both types of mutations involve changes from G to A or C to T nucleotides, and both occur in regions with similar recombination rate profiles (Figure S2), they should, on average, be subject to similar strengths of biased gene conversion. Comparing hominoid and monkeys, the substitutions involving CpG transitions are on average 1.07-fold higher in lineages leading from the hominoid-OWM ancestor to OWM than they are in lineages leading to hominoids. Considering the hominoid-NWM ancestor, substitutions are 1.19-fold higher in lineages leading to NWM than to hominoids (Figure 2). In contrast, when considering transitions at non-CpG G/C sites, there are on average 1.39-fold more substitutions from the hominoid-OWM ancestor to OWM lineages than to hominoid ones and 1.71-fold more from the hominoid-NWM ancestor to NWM than to hominoid lineages (Figure 2). Thus, CpG transition rates are more similar across species, as observed in comparisons of smaller datasets of primates and mammals (7, 31). These results are robust to the choice of species of OWM and apes used, e.g., using gorilla instead of chimpanzee or gibbon instead of orangutan yields similar differences (Figure S3, S4).

**Figure 2:**
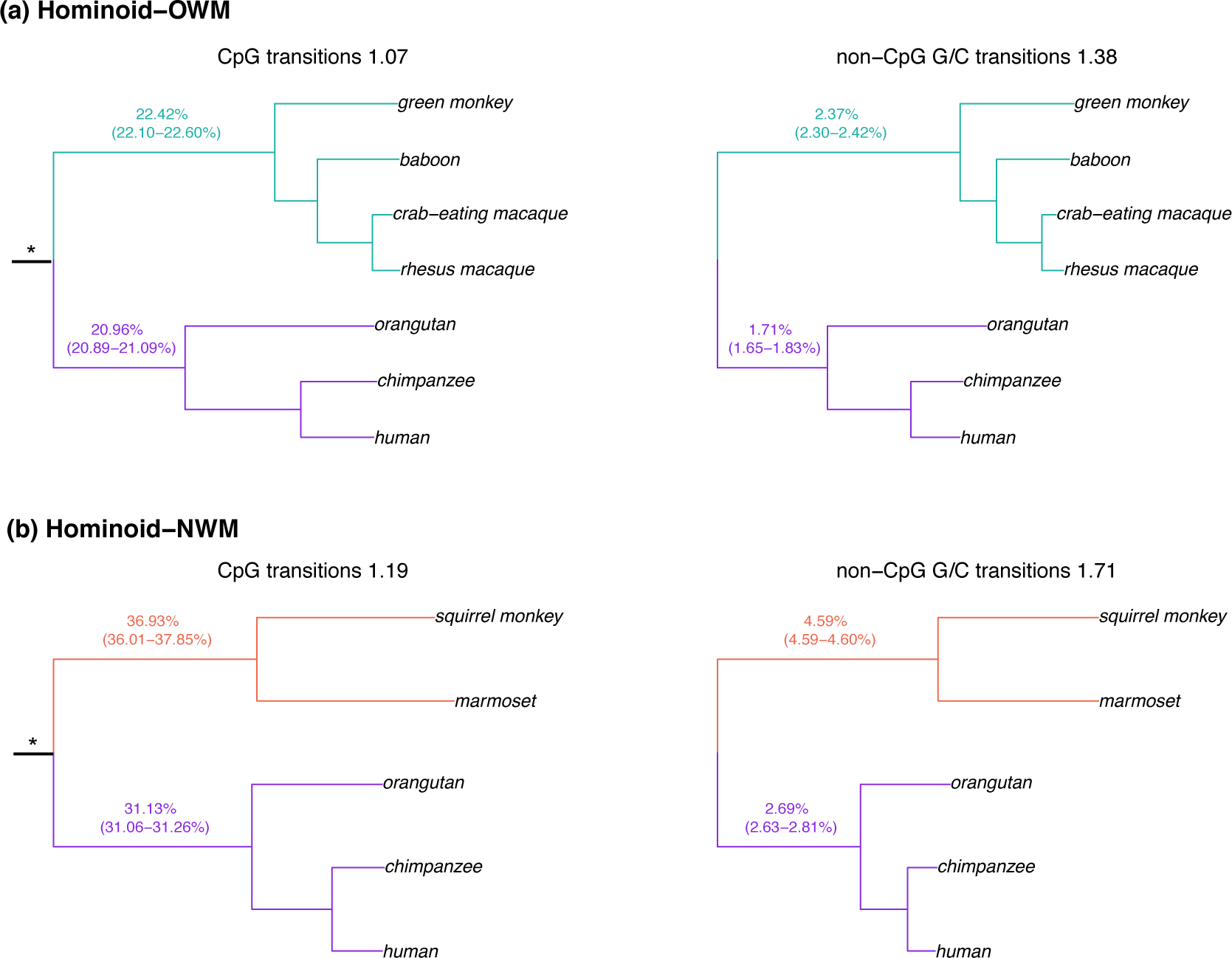
Comparison of substitution rates in hominoids and monkeys.

For transitions from CpG and non-CpG G/C sites, we estimate the total branch length from either (a) the hominoid-OWM ancestor to each leaf, or (b) the hominoid-NWM ancestor to each leaf. The mean length for monkeys (OWM, NWM) and hominoids are shown, along with range within each group. Branches from root-hominoids are shown in purple, from root-OWM in green and from root-NWM in orange. The symbol * indicates the hominoid-monkey (either OWM or NWM) ancestor used as root.

We then consider different substitution types in more detail, focusing on eight types: transitions and transversions occurring at either ancestrally A or T (referred to as A/T), ancestrally G or C (G/C), CpG and non-CpG G/C, again excluding CGI. As a measure of variation among species, we use the variance of the normalized root to leaf distances across all remaining nine species (see Methods), which is expected to be 0 if substitution rates are all identical. In general, transversions are more variable than transitions, with the largest variance (0.065) observed at A/T transversions (Figure 3a). In turn, the variance is lowest for CpG transitions outside of annotated CGI (0.005), as observed previously in comparisons of smaller datasets of 19 mammals (1.7 Mb) (7) and 9 primates (0.15 Mb) (60). Interestingly, transitions at CpG sites *inside* CGI have a greater variance in substitution rates, and behave similarly to G/C transitions (Figure 3a). The difference in behavior of CpGs inside and outside of CGI is again consistent with the notion that when the source of mutation is primarily non-replicative, mutations depend primarily on absolute time, whereas when they have other sources that are dependent on the numbers of cell divisions and are more variable among species. If this interpretation is correct, an interesting implication is that germline methylation levels and spontaneous deamination rates have remained very similar across primate species.

**Figure 3:**
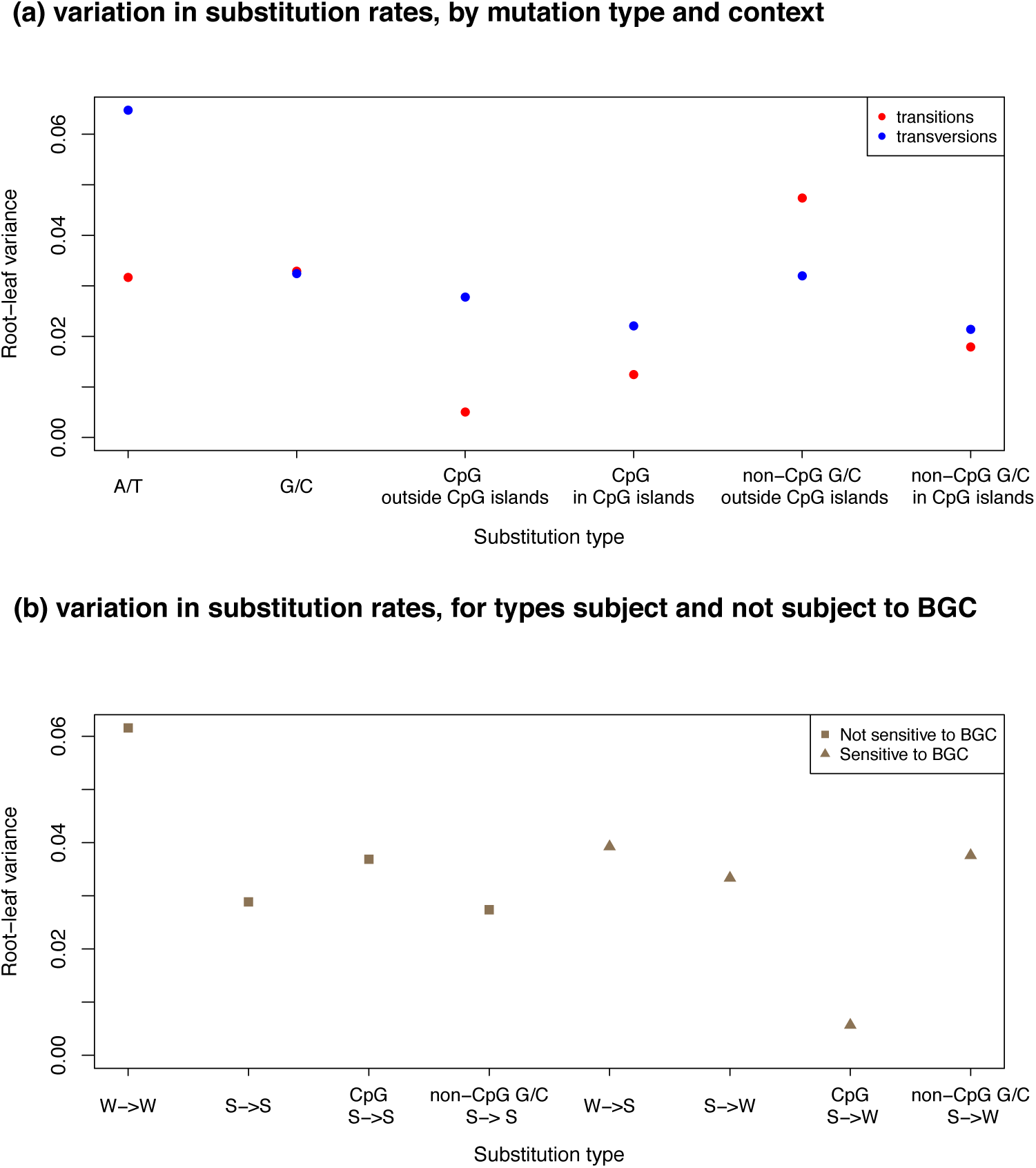
Variance among lineages for different substitution types. (a) For each ancestral state and each context shown on the x-axis, we estimate the total branch length from the root to each terminal leaf in the Multiz dataset as the inferred number of substitutions per site. We then compute the variance in the normalized root to leaf distance across nine primate species (human, chimpanzee, orangutan, rhesus macaque, crab-eating macaque, baboon, green monkey, squirrel monkey and marmoset). (b) For each substitution type (strong (S; G/C) and weak (W; A/T)) in different substitution contexts shown on the x-axis, we estimate the total branch length from the root to each terminal leaf in the Multiz dataset, and compute the variance in the root-leaf distance across the nine primates used in (a).

**Figure 4:**
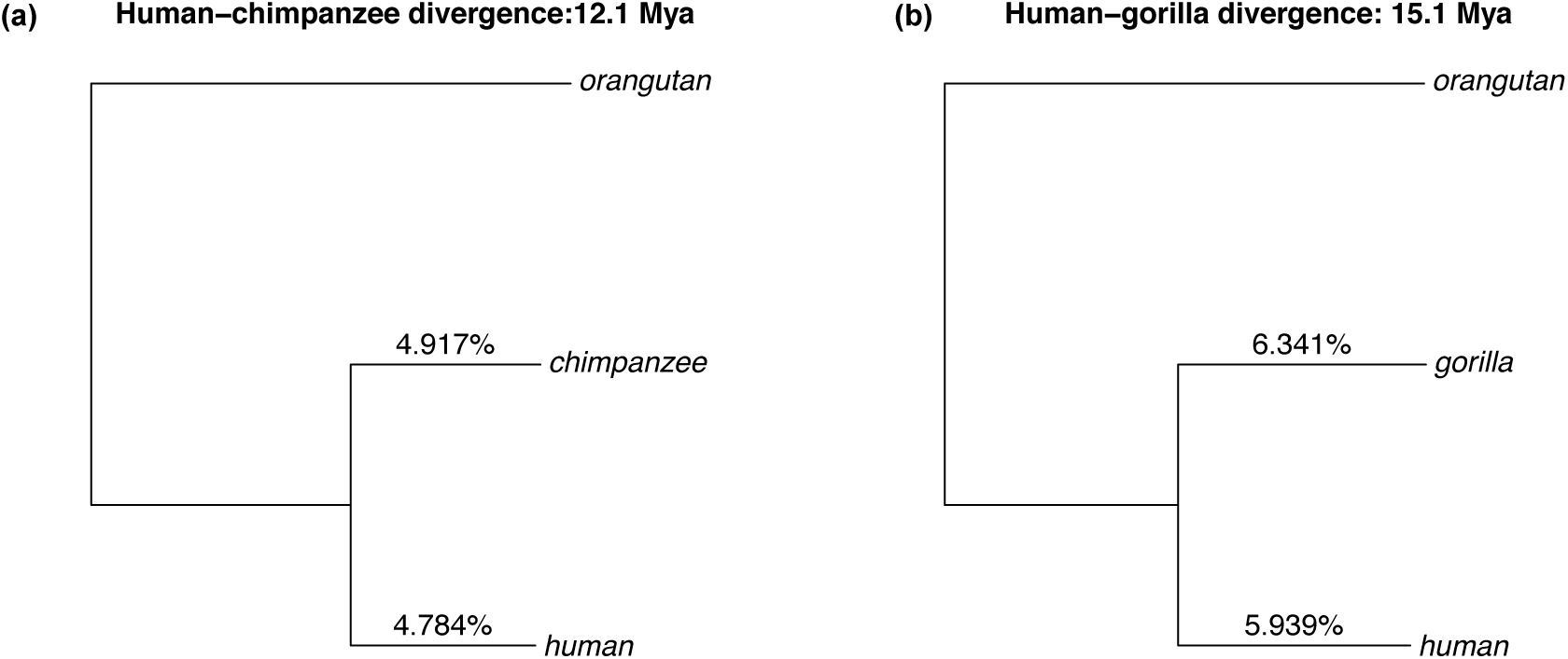
Revised estimates for divergence of hominines. We estimate the autosomal substitution rates for transitions at CpG sites by applying Phylofit to the high coverage pairwise alignment of human, chimpanzee and human and gorilla mapped to the orangutan reference genome. Divergence time is estimated from rates of *de novo* CpG transitions obtained in human pedigrees (see Methods for details).

Patterns of substitutions may also vary across species due to effects of biased gene conversion, notably because of differences in effective population sizes (26). To examine this possibility, we compare the variance of the normalized root to leaf distances for substitutions that are subject to BGC (such as W-> S or S->W) and those that should not be affected by BGC (such as W->W and S->S). If the strength of BGC varies across primates, we expect larger variance across species at W->S and S->W substitutions. Instead, there is no significant difference in the estimates of variances across the two classes of substitutions (Figure 3b; p = 0.5, based on a permutation test). While this finding seems puzzling given the three to four-fold difference in effective population size of these species (28), it is consistent with results of Do et al. (2015), who found no significant difference in the extent of biased gene conversion across diverse groups of West African and non-African human populations that differ up to two-fold in their effective population sizes (61). If the strength of biased gene conversion at a site is typically very weak, both findings could reflect lack of power.

Given the importance of a steady molecular clock for dating events in human evolution, we next focus specifically on hominines. In these comparisons, subtle differences in sequence quality, coverage, or the extent of mapping artifacts can lead to misleading evidence for variation in substitution rates across species. To minimize these effects, we generated pairwise sequence alignments for high coverage (˜30x) genomes of human and chimpanzee and human and gorilla. These pairs of genomes were mapped to the orangutan reference genome (which should be equidistant to all three species, assuming no differences in substitution rates among species), matching the alignment and variant calling pipeline for all three species (see Methods). In this way, and after removing missing data, non-neutral sites and CGI, we obtain ˜1.03 Gb for human-chimpanzee and ˜1.02 Gb of human-gorilla whole genome sequence alignments.

Despite the differences in generation time and onset of puberty among extant chimpanzees and humans, rates of evolution on the two lineages are very similar: 0.621% and 0.633% respectively. This difference of 1.9% is however highly statistically significant under the assumption of no systematic errors (p < 10^-20^). When we consider the substitution rates at different mutation types, there is somewhat more pronounced differences for some types of substitutions, in inconsistent directions. For example, when comparing chimpanzee to human branches for substitutions involving transversions from CpG sites, the difference is 0.91-fold, whereas it is 1.07-fold for transversions at A/T sites (Figure S9). Comparing human and gorilla lineages, differences in substitution rates are more pronounced: the gorilla branch (0.824%) is longer than that the human (0.773%) branch by an average of 6.6% (p < 10^-20^). Again, different types of substitutions show distinct patterns, ranging between 0.96-fold at CpG transversions on the gorilla versus human branches to 1.10-fold for A/T transitions (Figure S10).

To check the reliability of these inferences, we also estimate the substitution rates using a second, newly developed maximum-likelihood based method (see Methods). While the absolute values of the substitution rates differ between the two methods, possibly due to methodological differences in calling ancestral states and assumptions about stationarity, the ratio of substitution rates between humans and chimpanzees (Figure S9, S11) or between humans and gorillas (Figure S10, S12) is almost identical. While these results match those obtained by Elango et al. (2005) based on 75 Mb of human-chimpanzee sequence, they are only about two-thirds of what was reported as the difference between humans and gorilla, based on 2 Mb of sequence data (32). Since our study is able to take advantage of a much larger dataset, accounts for differences in coverage and mapping among reference genomes and considers only putatively neutral sites, we surmise that the earlier estimate of the extent of substitution rate variation among human and gorilla was somewhat too high.

Importantly, these observations highlight that the mutation spectra, and not just the yearly mutation rate, are changing across primates. Notably, while the rate of substitutions involving CpG transitions is relatively stable across species, those of other substitution types vary substantially. As a result, the proportion of substitutions that consist in CpGs versus other mutation types differs among primates (Table S3). Since a transition to transversion ratio of 2.2, as found in humans, is often used as an indication of high quality genotypes (62), a practical implication is that this criterion cannot be reliably exported to other primates, or beyond. More fundamentally, our findings underscore that the mutation spectrum appears to have changed over the course of primate evolution. In this regard, it mirrors observations from even shorter time scales: for example, the recent report that transitions from 5’-TCC-3’ → 5’-TTC-3’ occurred at a proportionally higher rate in Europeans compared to Asians and Africans since these populations split (63).

## Discussion

Evolutionary rates are faster in NWM compared to OWM and in turn; rates in OWM are faster than in humans and apes. These findings support the hominoid rate slowdown hypothesis (64, 65), indicating that since the split of hominoids and monkeys, per year mutation rates have decreased considerably. Moreover, the ordering of substitution rates is consistent with the generation time hypothesis, in that NWM have a substantially shorter generation time (g = ˜6 years (20)) than OWM (g = ˜11 years (66)), who in turn reproduce at younger ages than apes (g = ˜25 years). Within hominines, gorillas (g = ˜19 years (67)) have a faster yearly rate than humans (g = ˜29 years) and chimpanzees (g = ˜25 years (67)). To investigate if the association between generation time and substitution rates is significant after controlling for the underlying phylogeny, we perform independent contrast analysis (53). Specifically, we assume the underlying phylogeny based on CpG transition rates (effectively assuming these are strictly clock-like) and then estimate the correlation between generation times and non-CpG substitution rates, controlling for the shared phylogenetic history. While we have few species available for the analysis, the results are not significant (r = 0.17, p=0.7; see Methods), so the causal relationship remains to be established.

As an alternative approach is to ask whether differences in generation times and other life history traits can plausibly explain the variation in per year mutation rate. To this end, we consider a simple model for mutations that are replicative in origin (or non-replicative but well repaired (25)). Across primates, the average time between puberty and reproduction is positively correlated with age of onset of puberty in males (r = 0.74, p = 0.01, using Spearman’s correlation test), and the rate of spermatogenesis (measured by estimating the seminiferous epithelium cycle length (SECL)) is positively correlated with generation time by (r = 0.90, p = 0.002) (Table S4). We therefore vary the generation time, age of onset of puberty and SECL for each lineage, relying on values estimated for extant humans, chimpanzees and OWM (Table S4). On that basis, we predict that yearly mutation rates in humans and chimpanzees should differ by ˜30%, and hominoids and OWM should differ by ˜66%. Thus, if anything, differences in life history traits predict even more variation in substitution rates than is observed.

Of course, these species may have had similar life histories throughout much of their evolutionary past, such that the use of life-history traits estimated in extant species exacerbates expected differences. For instance, fossil evidence suggests that the age of puberty on the human lineage may have only recently increased, and was lower in *Homo erectus* and Neanderthals (30, 68). If we change the age of onset of puberty in humans to 9 years, the difference in rates between humans and chimpanzees is only expected to be One implication, then, of finding such similar substitution rates in humans and chimpanzees is that their life histories are likely to have been pretty similar for much of their evolutionary history.

That substitution rates should and do vary with life history traits highlights the challenges of using the molecular clock for dating evolutionary events, even within hominines. All the more so when there are substantial differences in the generation time between males and females (59). One way to overcome this difficulty is to explicitly model the changes in life-history traits over the course of primate evolution and study their effect on substitution rates (59). Notably, Amster and Sella (2015) show that accounting for variation in generation time, age of onset of puberty and rate of spermatogenesis in apes helps to reconcile the split times estimated based on molecular and fossil evidence (59). This approach, however, requires knowledge of life-history traits in extant and ancestral populations, which is hard to obtain for many species.

An alternative may be to focus on mutation types that are much less sensitive to life-history traits, such as CpG transitions outside CGIs. Even for this mutation type, the variance in substitution rates across species is non-zero, possibly because a subset of these mutations occurs at unmethylated sites or through replication errors, or because repair is not completely inefficient. Nonetheless, CpG transitions accumulate in a quasi-clock like manner and appear to be least affected by life history differences across species. Moreover, in humans, they contribute almost a fifth of all *de novo* mutations (12).

With these considerations in mind, we re-estimate the divergence and split times of human, chimpanzees and gorillas using substitution rates estimated only at CpG transitions. Assuming the per year mutation rate for CpG transitions obtained in (12) (see Methods), we estimate that humans diverged from chimpanzees ˜12.1 million years ago (Mya) and from gorillas ˜15.1 Mya. Assuming further that the effective population size of the human-ape ancestor was five times the current population size (as estimated by (37, 56)), the human-chimpanzee split time is approximately 7.9 Mya and human-gorilla split time of 10.8 Mya. Reassuringly, these estimates are similar to those obtained by explicitly modeling the dependence of replicative mutations on life history traits in hominines (59). Moreover, these estimates are in broad agreement with evidence from the fossil record, which suggests a human-chimpanzee split time of 6-10 Mya and human-gorilla split time of 7-12 Mya (69-72). Thus, perhaps there is no real discrepancy between phylogenetic and pedigree based estimates of mutation rates, once the impact of life history traits on mutation rates is taken into account (59).

## Acknowledgments

We thank Guy Amster, Ziyue Gao, Minyoung Wyman, David Pilbeam and Guy Sella for helpful discussions. We thank Nick Patterson and Heng Li for technical advice for mapping and alignment of high coverage genomes, Melissa Hubisz and Adam Siepel for advice on running Phylofit. PM was supported by the National Institutes of Health (NIH) under Ruth L. Kirschstein National Research Service Award F32 GM115006-01. CEGA was supported by a Science Without Borders fellowship from CNPq - Conselho Nacional de Desenvolvimento Científico e Tecnológico - Brazil (PDE 201145/2015-4).

## Supplementary material

### Note S1: Generating high coverage data for human genome

For our high coverage analysis, we sequenced one individual of European ancestry. This individual provided informed consent for participation in the study. The project was approved by the Institutional Review Boards (IRBs) at The University of Chicago and Columbia University.

Genomic DNA was extracted from blood and one library was constructed using Illumina-recommended protocols. Briefly, 1μg of DNA was extracted and sheared into fragments using sonication. The resulting fragments were end repaired, adenosine overhangs were added and adaptors were ligated. Gel electrophoresis was performed to select libraries with insert sizes of approximately 350 bp in size, which were amplified using quantitative PCR. The resulting libraries were sequenced using Illumina HiSeq 2000 to generate paired-end reads. We generated ˜49 Gb of sequencing data (˜30x coverage). Mapping and alignment were done using samtools as described in the main text.

Sequence data will be available upon publication from dbGap from the following link: https://www.ncbi.nlm.nih.gov/projects/gap/cgi-bin/study.cgi7study_id=phs000599.v2.p1

### Note S2: Analysis of EPO dataset

To test the robustness of our inferences, we repeat the analysis with the EPO dataset containing seven primates (human, chimpanzee, gorilla, orangutan, rhesus macaque, baboon and marmoset). Due to concerns of incomplete lineage sorting between chimpanzee/gorilla (51), we exclude gorilla from further analysis. After filtering putatively selected sites and removing missing data, we analyze approximately 745 Mb of whole genome sequence alignment. To allow for direct comparison with the Multiz dataset, we repeat our main analysis with the same smaller subset of species available for the EPO dataset. Due to challenges in accurately reconstructing the ancestral state for outgroup species, here marmoset, substitution rates in NWM could be underestimated and hence we do not include comparisons of hominoids and NWM for this dataset.

We apply Phylofit to estimate the substitution rates across all species (Figure S5) and find that substitution rates on lineages leading from the hominoid-OWM ancestor to hominoids are on average 2.85% (range 2.81-2.88% across species) whereas rates on lineages leading to OWM are on average 3.59% (3.589-3.593%), 1.26-fold higher. These estimates are lower than results reported in the main text, as we are using a smaller subset of species. Similar estimates are obtained for the smaller subset of species with the Multiz dataset where the substitution rates are 1.28-fold faster in OWM compared to hominoids. We also repeat the main analyses shown in Figure 2 and 3 with the smaller subset of species in the EPO and Multiz dataset (Figure S6-S8).

**Figure S1:**
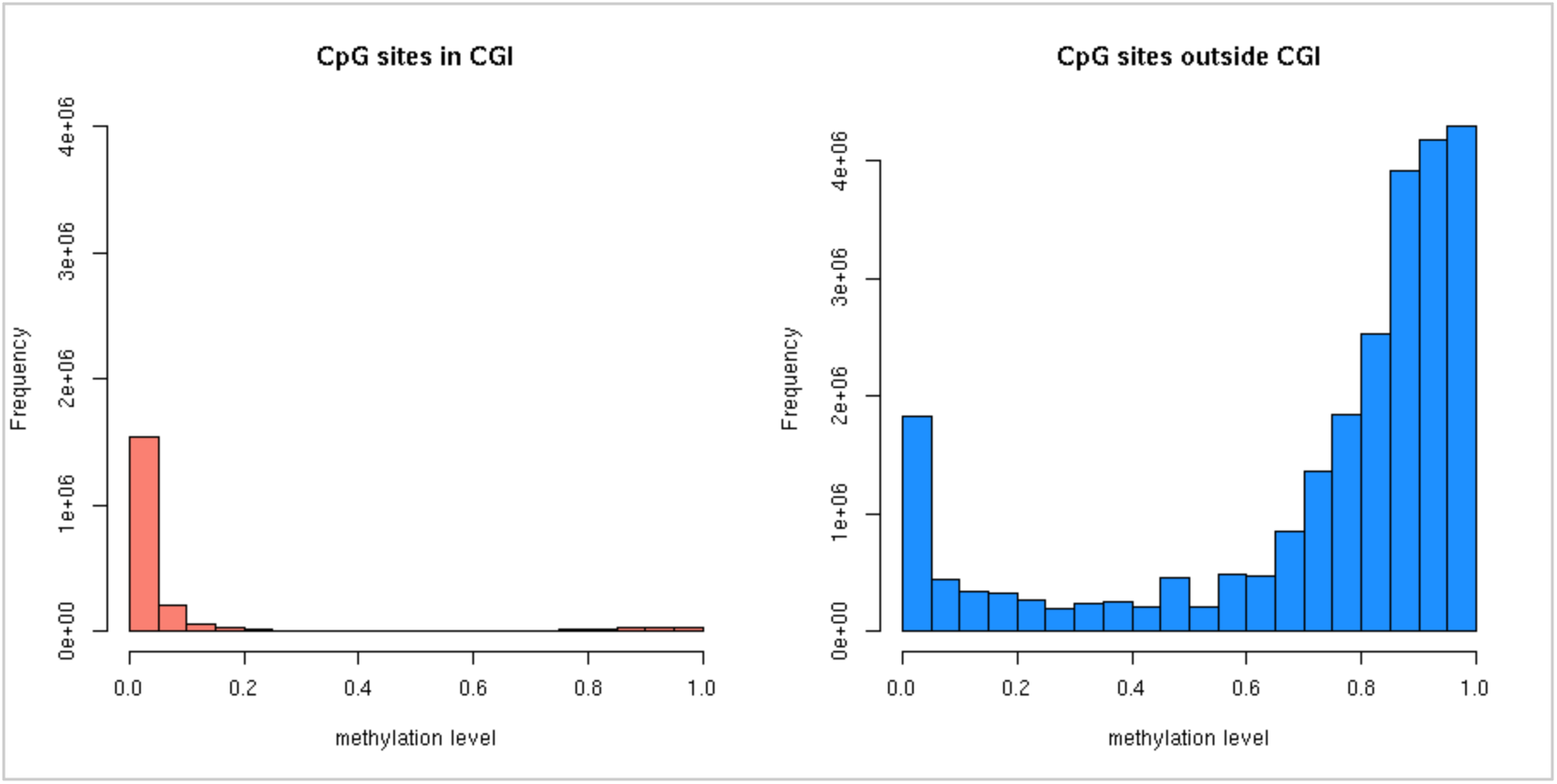
Sperm methylation profiles of CpG sites. We plot the distribution of methylation levels at CpG sites inside and outside of annotated CGI. The methylation profiles in human sperm were taken from (47).

**Figure S2:**
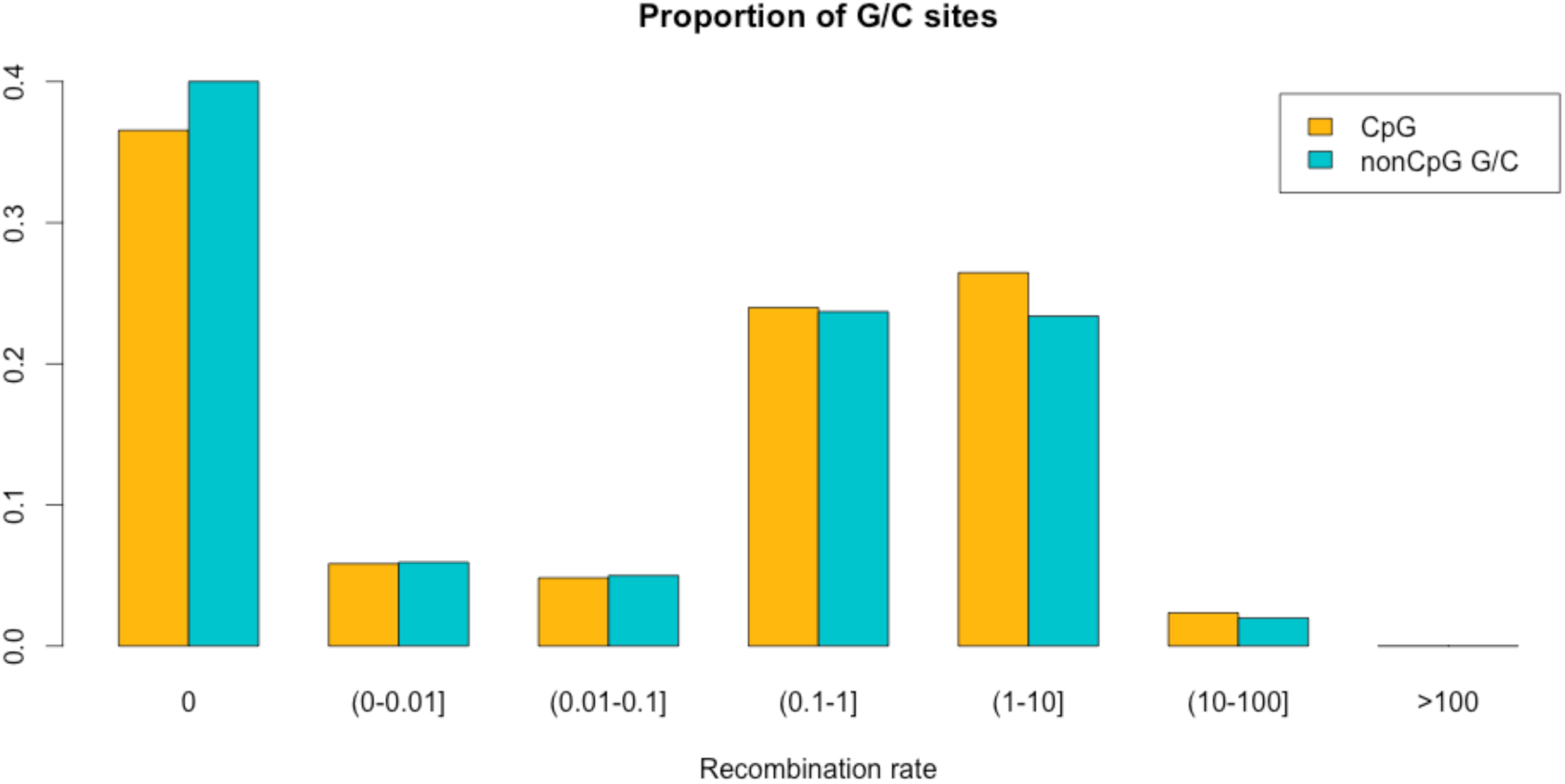
Distribution of CpG and non-CpG G/C sites across the human genome. We plot the proportion of CpG and non-CpG G/C sites in the human genome as a function of recombination rate. After filtering non-neutral sites and CGI (see Methods) in the Multiz dataset, we estimate the overall proportion of CpG and non-CpG G/C sites as 1.60% and 37.9% respectively. Crossover rates were obtained from the deCODE genetic map, UCSC genome browser track “deCODE Recombination maps: Sex avg” (73) originally calculated in 10 kb bins and standardized to have an average rate of 1.

**Figure S3:**
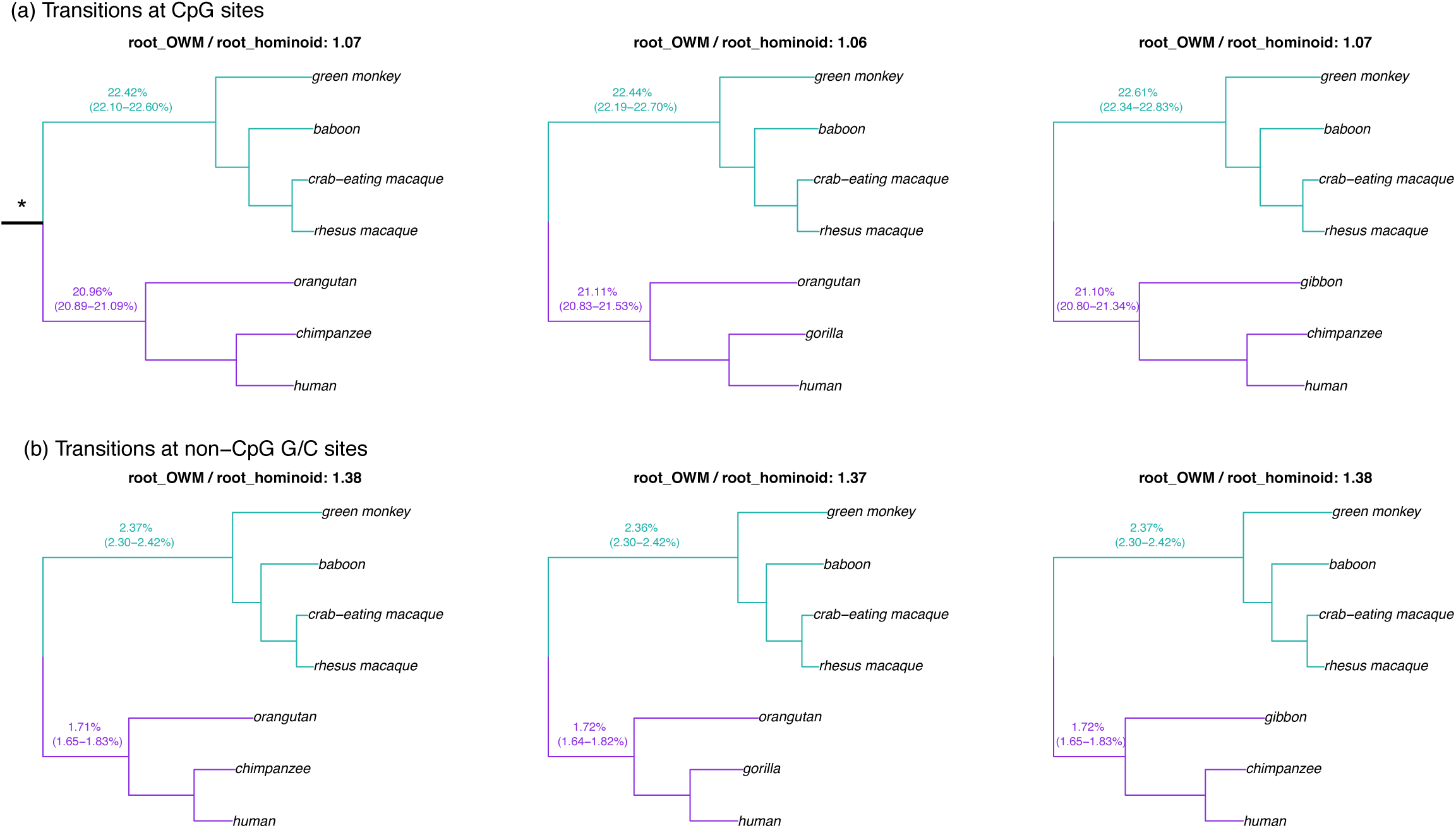
Comparison of substitution rates in hominoids and OWM using alternate topologies. Due to concerns about the possible effects of incomplete lineage sorting, we analyze gorilla and chimpanzees and gibbon and orangutans separately. Each sub-figure shows a different set of species and substitution type (CpG / non-CpG G/C sites). For each topology, we estimate the total branch length from the hominoid-OWM ancestor to each leaf. The mean lengths for OWMs and hominoids are shown, along with range within each group. Branches from root to hominoids are shown in purple and from root to OWMs are shown in green. The symbol * indicates the hominoid-OWM root from which branch lengths were estimated. The ratio of the average substitution rate leading to an OWM to the average rate leading to a hominoid is shown as the title for each sub-figure.

**Figure S4:**
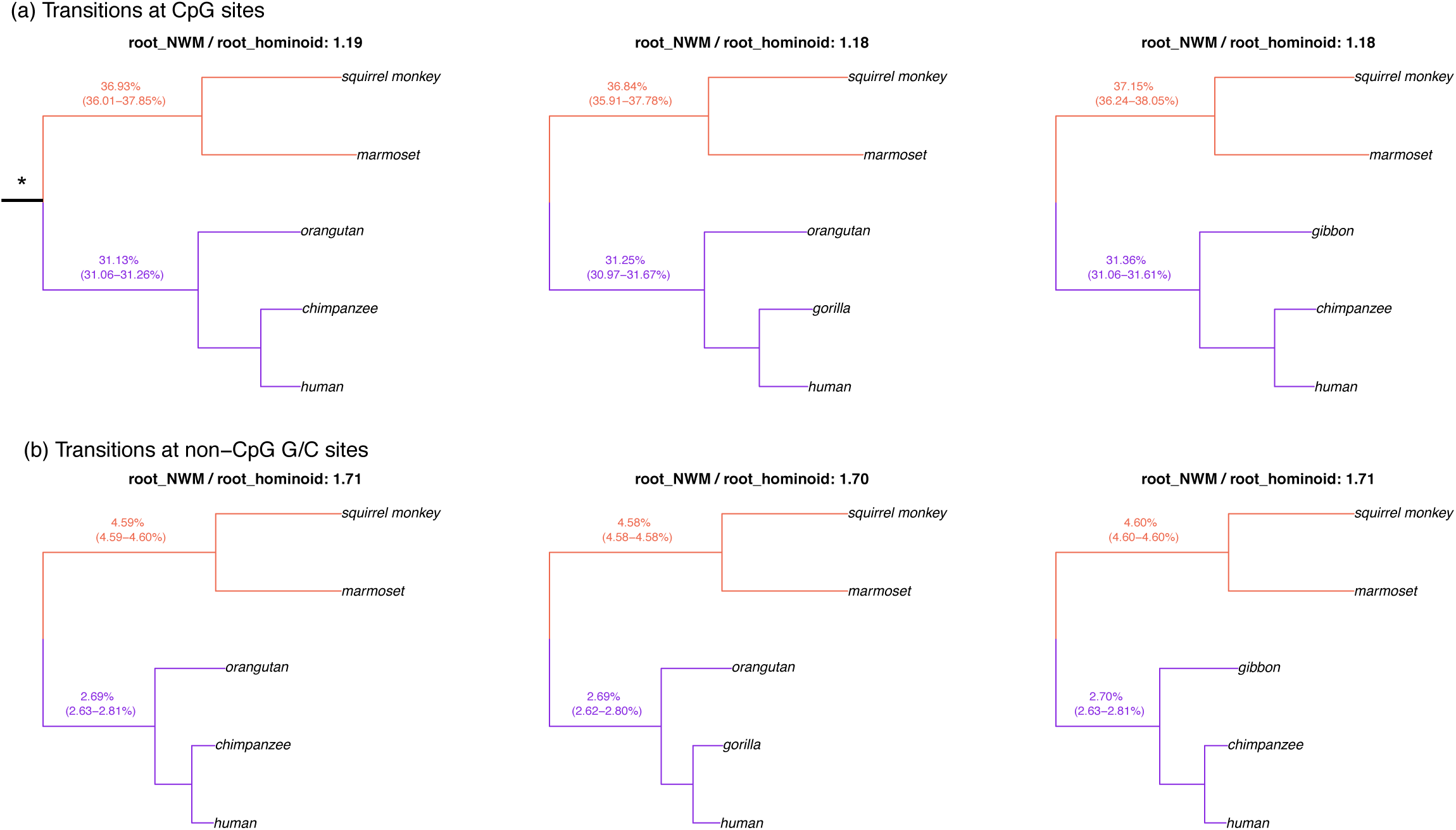
Comparison of substitution rates in hominoids and NWM using alternate topologies. Due to concerns about the possible effects of incomplete lineage sorting, we analyze gorilla and chimpanzees and gibbon and orangutans separately. Each sub-figure shows a different set of species and substitution type (CpG / non-CpG G/C sites). For each topology, we estimate the total branch length from the hominoid-NWM ancestor to each leaf. The mean lengths for NWMs and hominoids are shown, along with range within each group. Branches from root to hominoids are shown in purple and from root to NWMs are shown in green. The symbol * indicates the hominoid-NWM root from which branch lengths were estimated. The ratio of the average substitution rate leading to an NWM to the average rate leading to a hominoid is shown as the title for each sub-figure.

**Figure S5:**
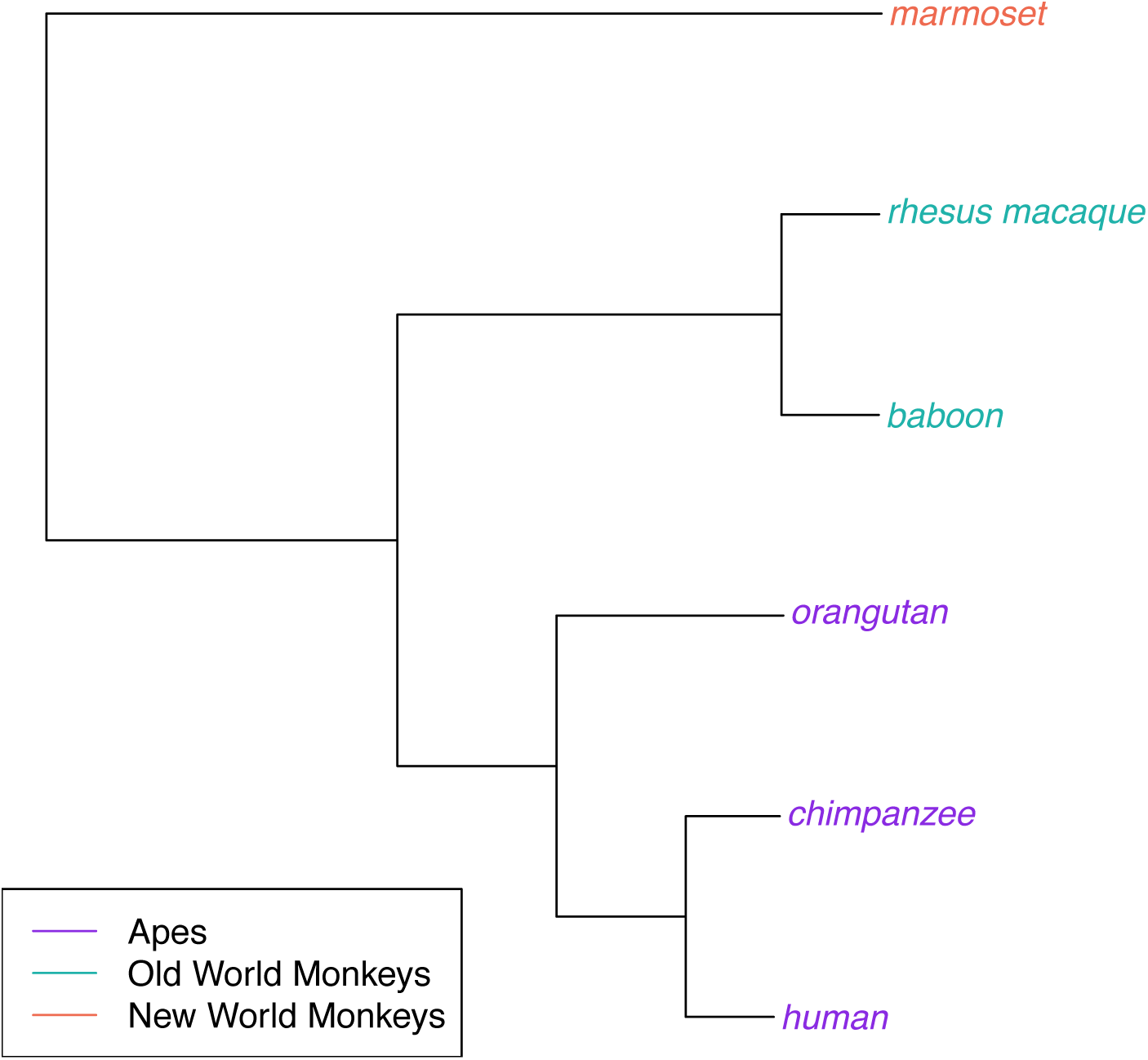
Phylogenetic tree for the six primates in EPO dataset. We estimated neutral substitution rates for six primates from the EPO dataset using Phylofit (see Methods for details). Branch lengths reflect the expected number of neutral substitutions per site along each lineage. We excluded gorilla due to concerns of possible effects of incomplete lineage sorting on estimates of substitution rates.

**Figure S6:**
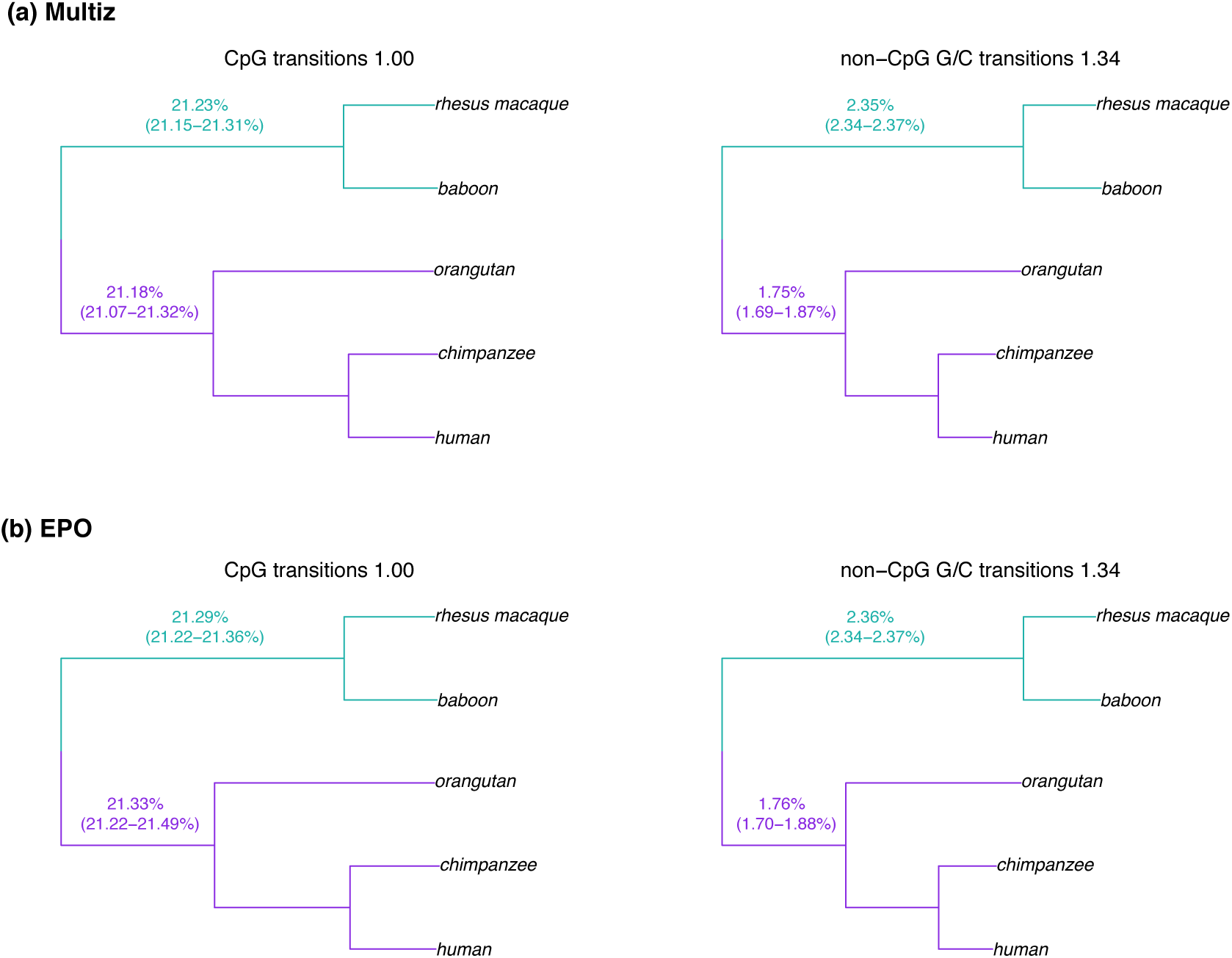
Comparison of substitution rates in hominoids and OWM using different datasets. For each dataset (Multiz or EPO), we estimate the total branch length from the hominoid-OWM ancestor to each leaf. The mean lengths for OWM and hominoids are shown, along with the range within each group. Branches from root to hominoids are shown in purple and from root to OWM are shown in green. The ratio of the average substitution rate on branches leading to OWMs to the average rate on branches leading to hominoids is shown as the title for each sub-figure along with the substitution context.

**Figure S7:**
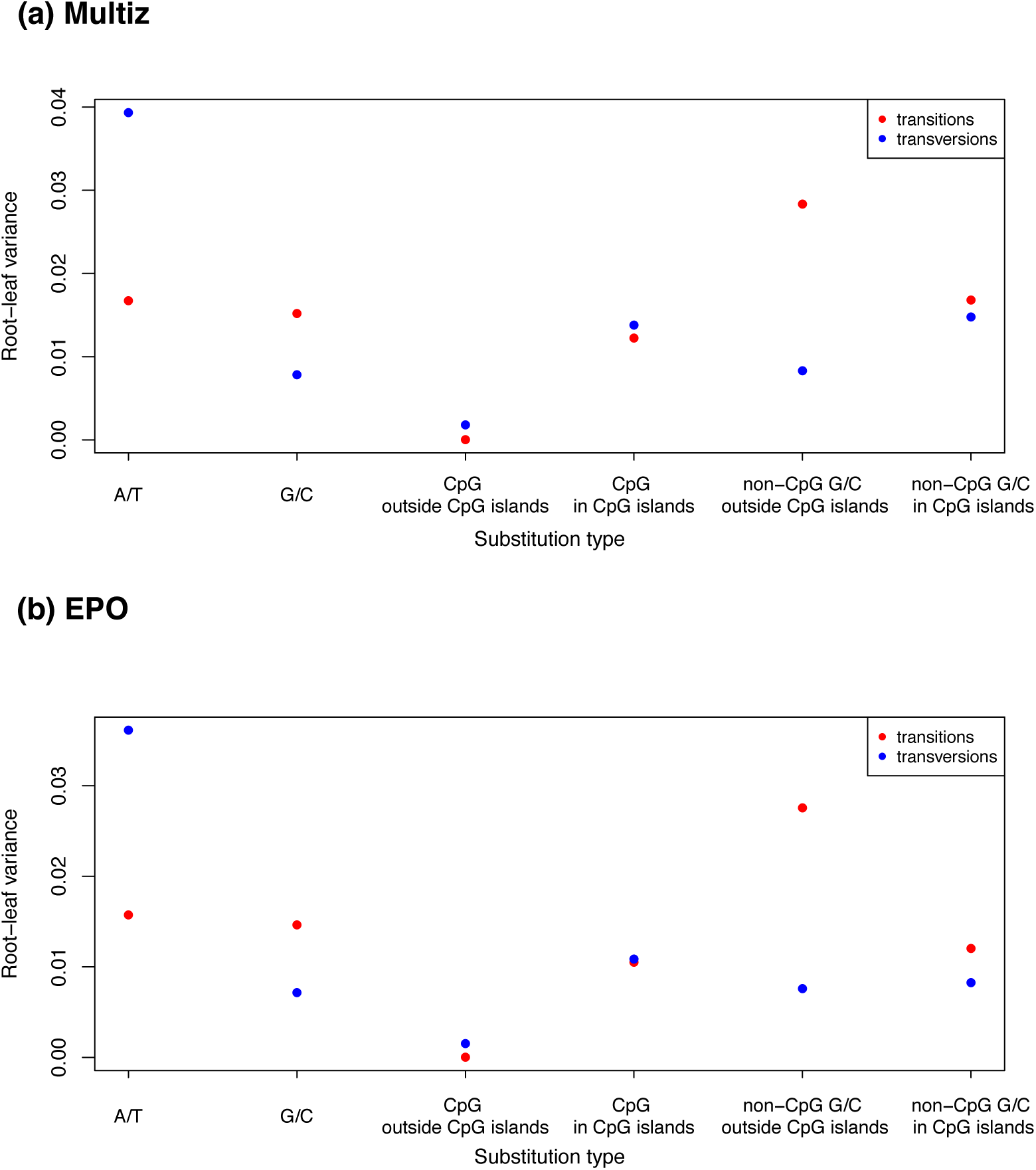
Variance among lineages for distinct substitution types, estimated from different datasets. For each ancestral state and each context shown on the x-axis, we estimate the total branch length from the root to each terminal leaf in the Multiz and EPO dataset as the inferred number of substitutions per site. We then compute the variance in the normalized root to leaf distance across five primates (human, chimpanzee, orangutan, rhesus macaque and baboon). This figure differs from Figure 2a, as it uses fewer species in the Multiz dataset to match the types of species (hominoids and OWM) available in the EPO dataset.

**Figure S8:**
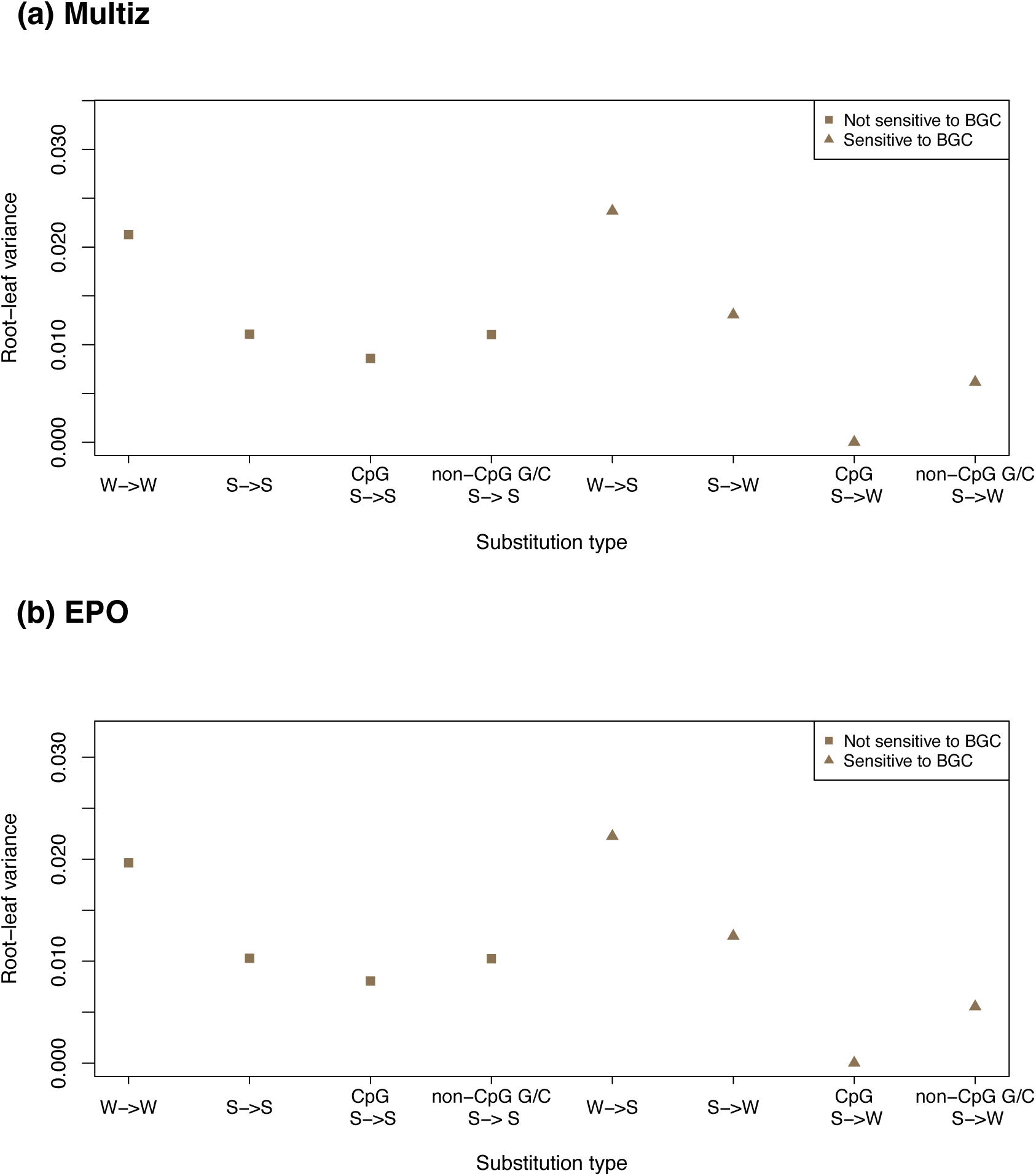
Effect of biased gene conversion across lineages estimated for different datasets. For each substitution type (strong (S; G/C) and weak (W; A/T)) and each ancestral context shown on the x-axis, we estimate the total branch length from the root to each terminal leaf in the Multiz and EPO dataset as the inferred number of substitutions per site. We then compute the variance in the normalized root to leaf distance across five primates in EPO (human, chimpanzee, orangutan, rhesus macaque and baboon). This figure differs from Figure 2b, as it uses fewer species in the Multiz dataset to match the types of species (hominoids and OWM) available in the EPO dataset.

**Figure S9:**
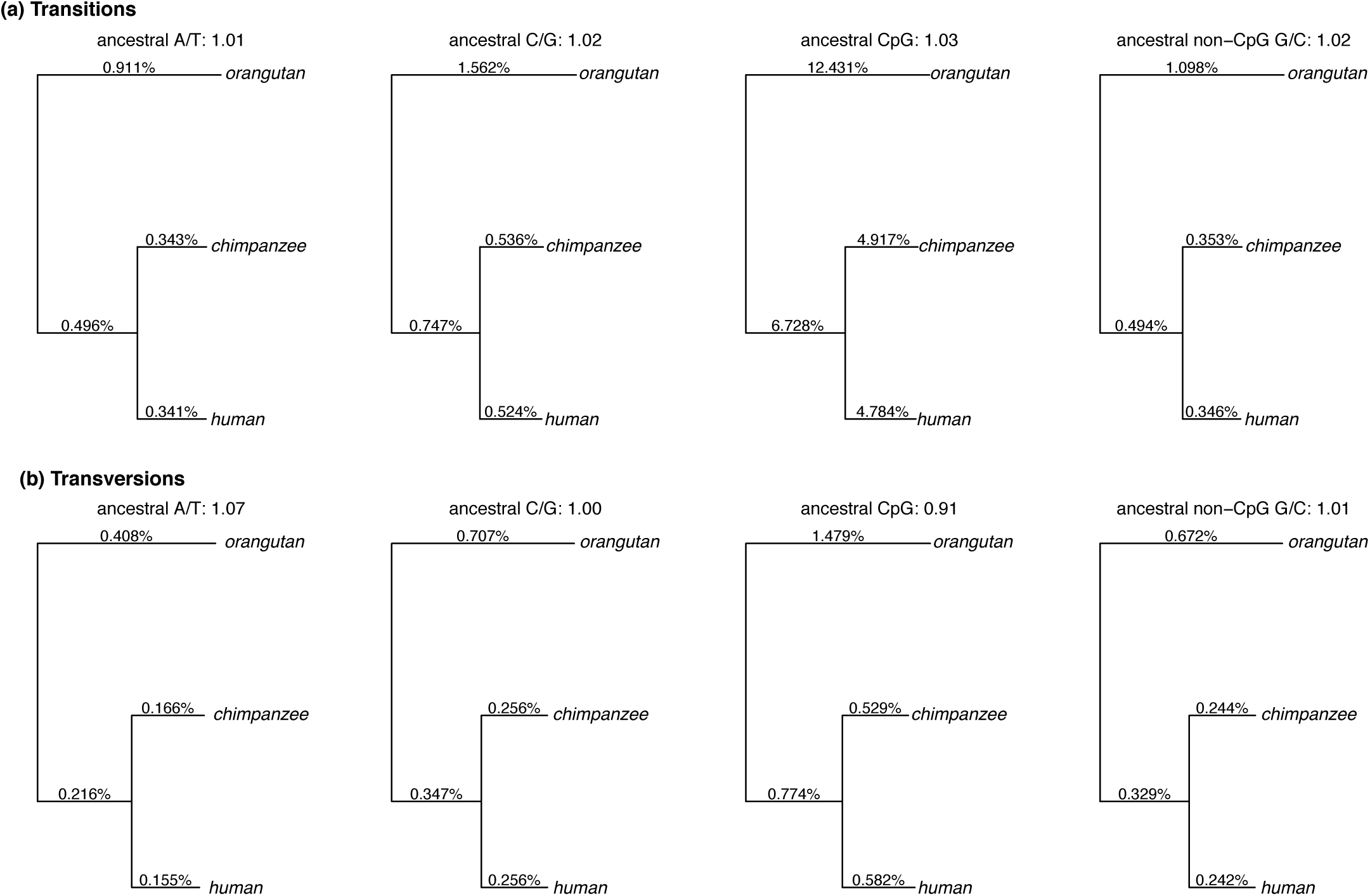
Comparison of substitution rates in humans and chimpanzees using Phylofit. For each substitution type, we estimate the autosomal substitution rate using the high coverage pairwise alignment of human and chimpanzee mapped to the orangutan reference genome. The ratio of substitution rate in chimpanzee to the substitution rate in human is shown as the title of each subfigure.

**Figure S10:**
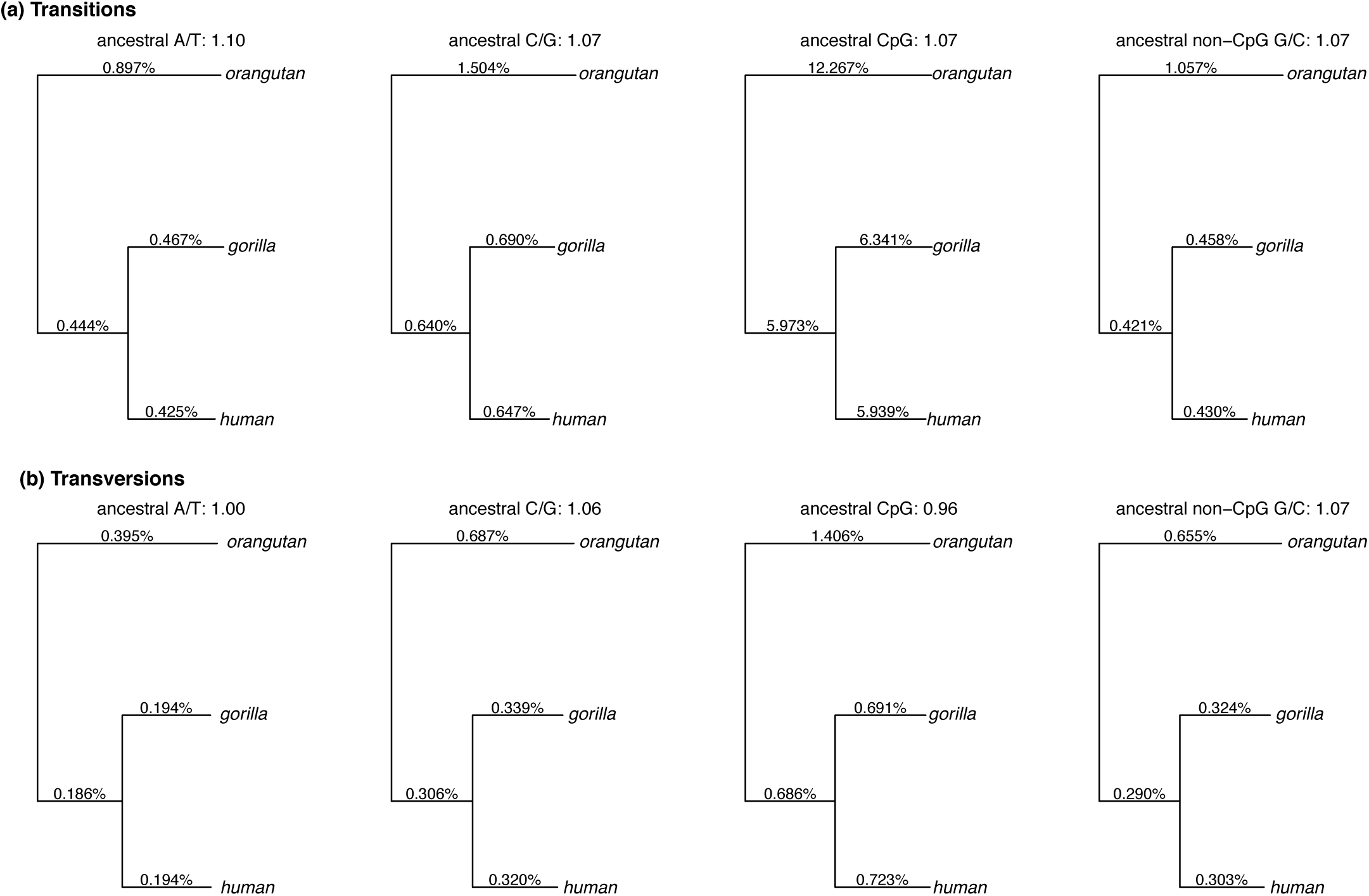
Comparison of substitution rates in humans and gorillas using Phylofit. For each substitution type, we estimate the autosomal substitution rate using the high coverage pairwise alignment of human and gorilla mapped to the orangutan reference genome. The ratio of substitution rate in gorilla to the substitution rate in human is shown as the title of each subfigure.

**Figure S11:**
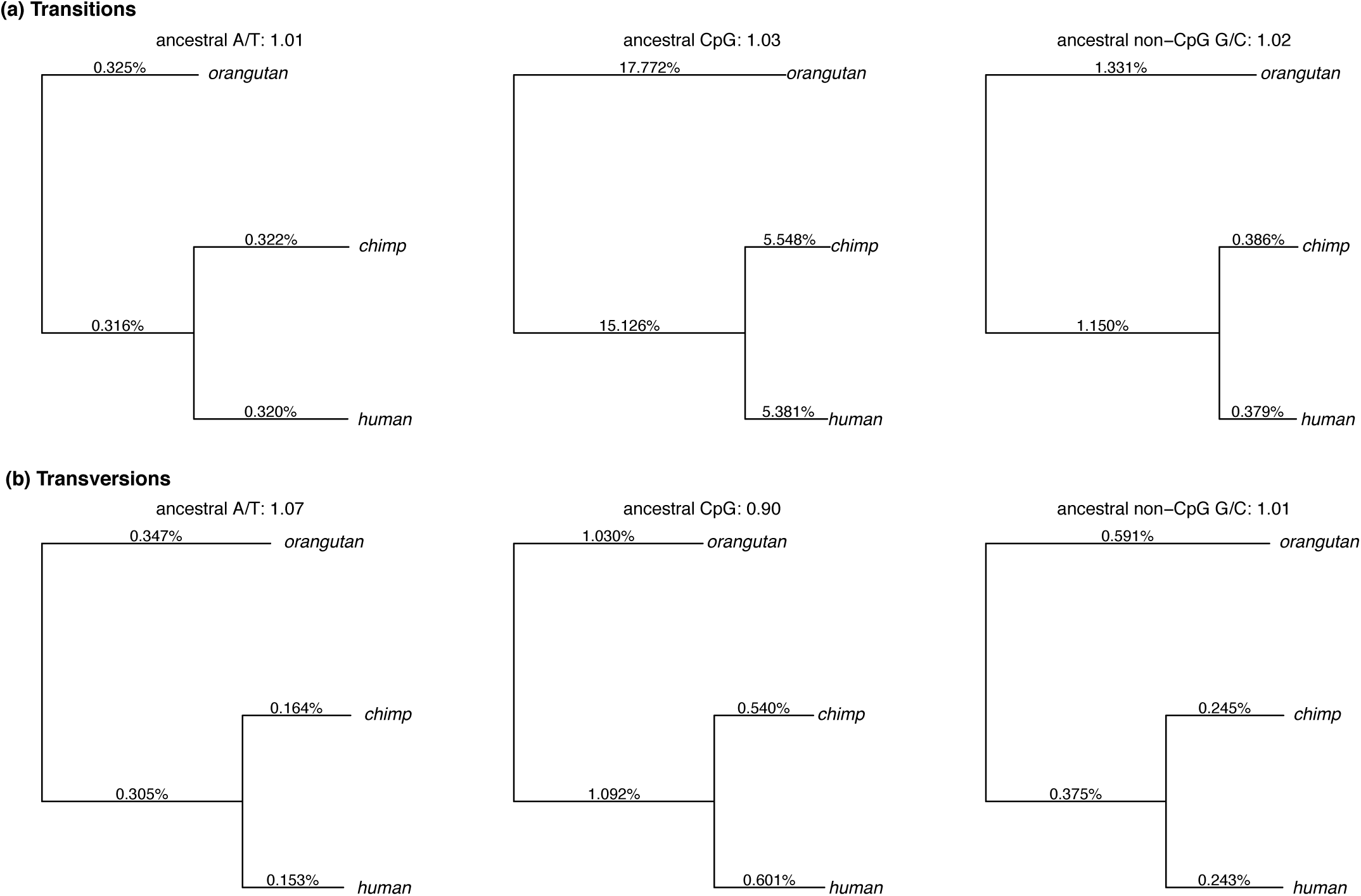
Comparison of substitution rates in humans and chimpanzees using the maximum likelihood approach. For each substitution type, we estimate the autosomal substitution rate using the high coverage pairwise alignment of human and chimpanzee mapped to the orangutan reference genome. The ratio of substitution rate in chimpanzee to the substitution rate in human is shown as the title of each subfigure. We note the maximum likelihood approach does not estimate the rates for ancestral C/G sites (that include CpG) and hence we do not show results for this context.

**Figure S12:**
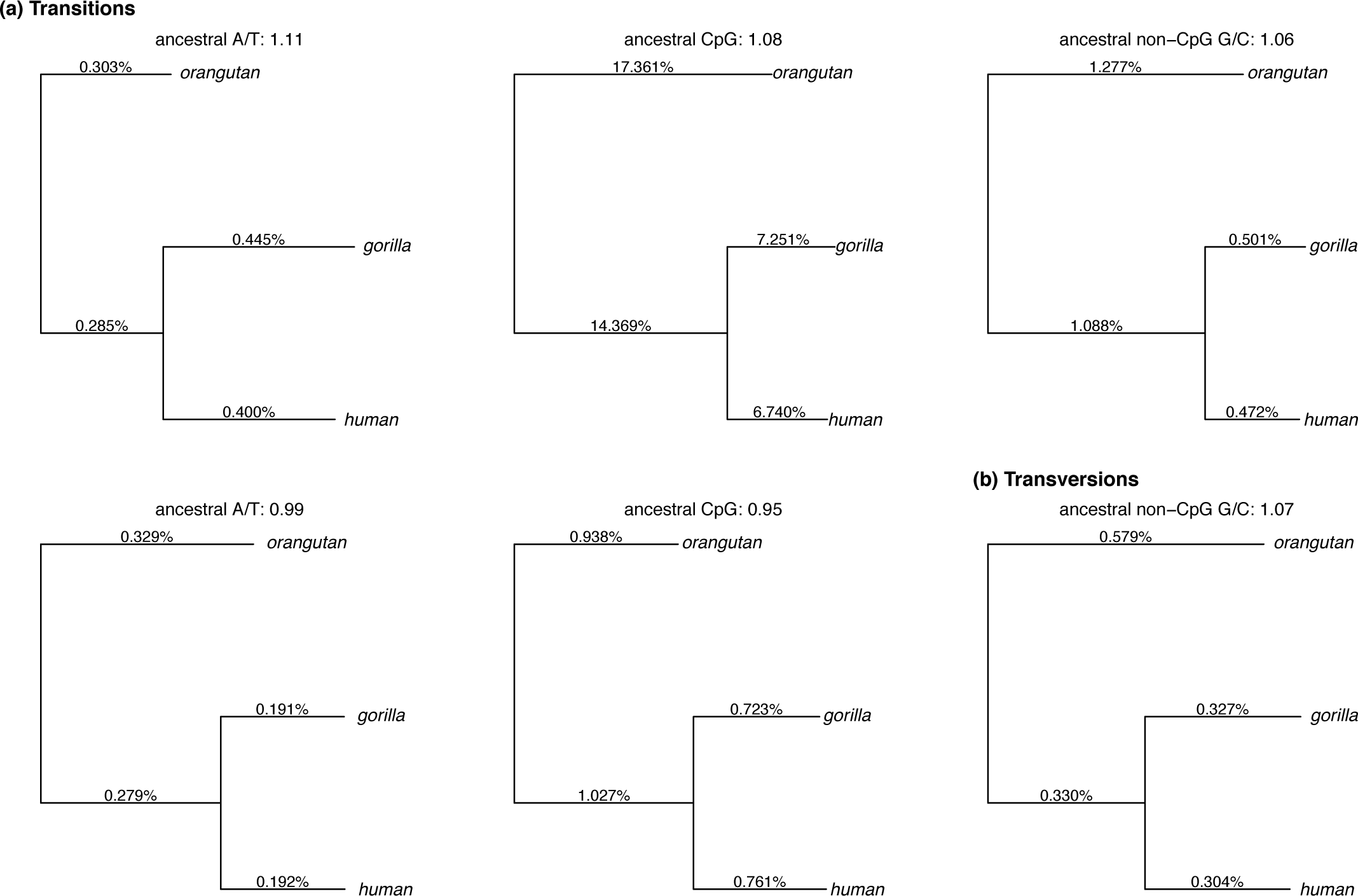
Comparison of substitution rates in humans and gorillas using the maximum likelihood approach. For each substitution type, we estimate the autosomal substitution rate using the high coverage pairwise alignment of human and gorilla mapped to the orangutan reference genome. The ratio of substitution rate in gorilla to the substitution rate in human is shown as the title of each subfigure. We note the maximum likelihood approach does not estimate the rates for ancestral C/G sites (that include CpG) and hence we do not show results for this context.

**Table S1.**
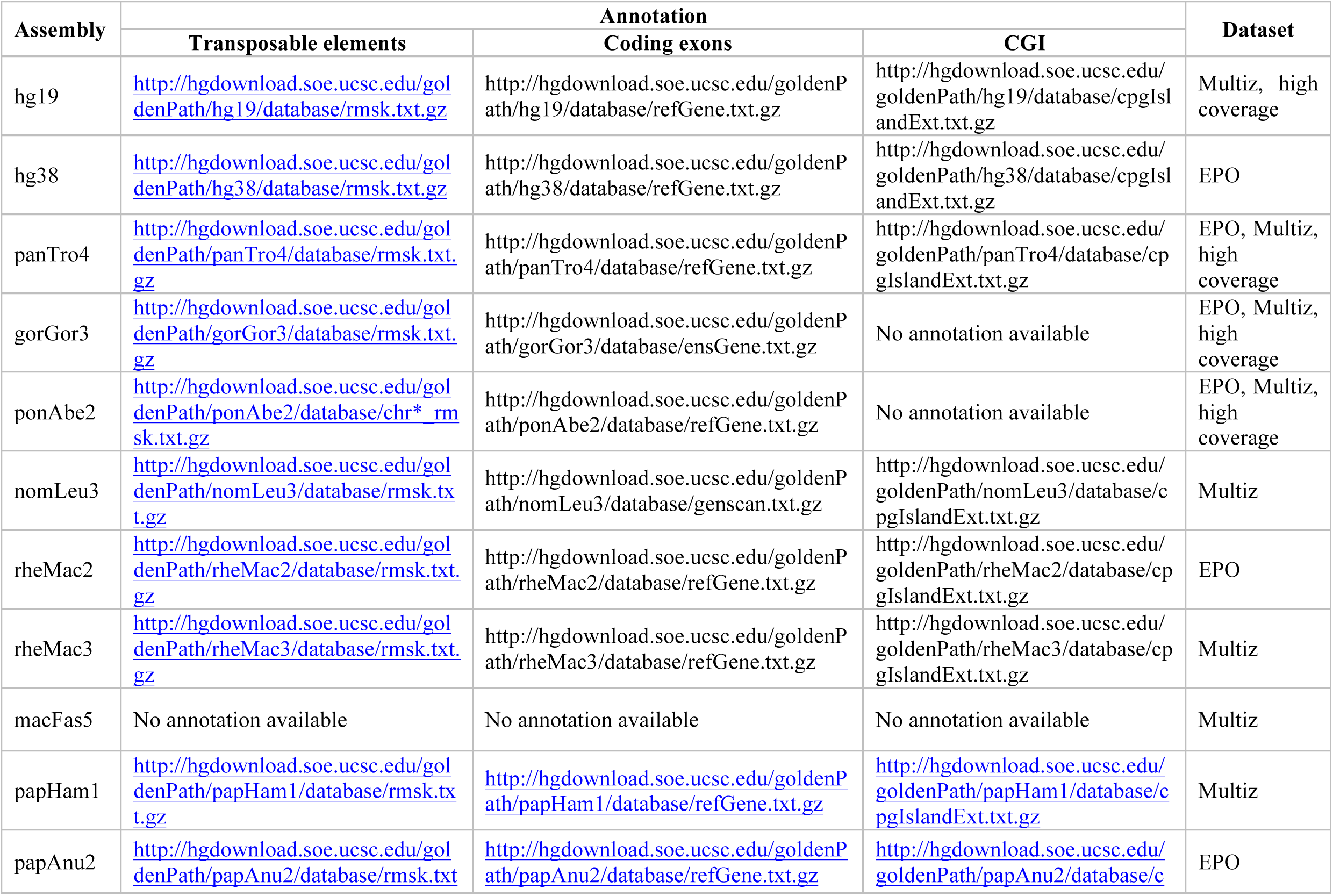
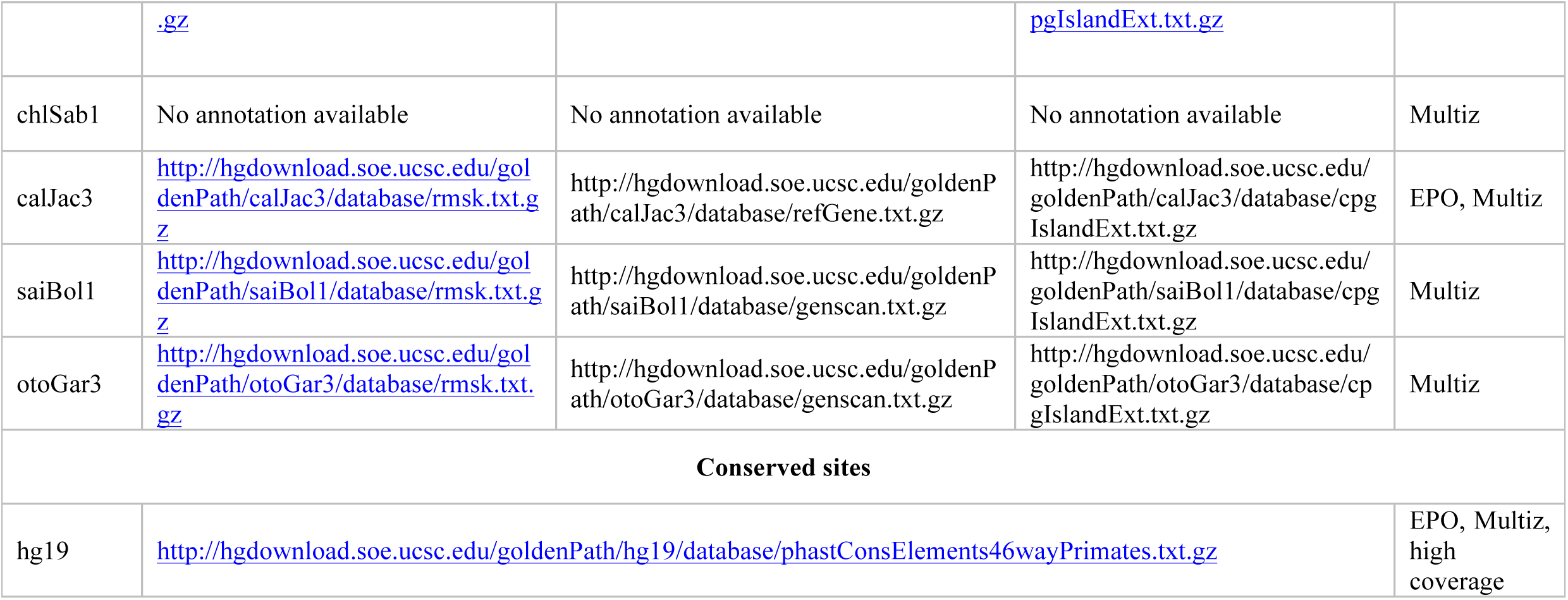
Online source of annotation for transposable elements, coding exons, CGI, and conserved sites.

**Table S2:** Life history traits in primates.

| **Species** | **Common name** | **SECL (in days)<sup>a</sup>** | **Generation time (in years)** | **Onset of puberty (in years)<sup>b</sup>** |
| --- | --- | --- | --- | --- |
| *Homo sapiens* | Human | 16 (74) | 29 (21) | 13.5 (75) |
| *Pan troglodytes* | Chimp | 14 (76) | 25 (67) | 8.5 (77) |
| *Gorilla gorilla* | Gorilla | n/a | 19 (67) | 7 (78) |
| *Pongo abelii* | Orangutan | n/a | 27* (79) | 6.5<sup>+</sup> (80) |
| *Macaca fascicularis* | Crab-eating macaque | 10.2 (81) | 11<sup>+</sup> (82) | 3.5 (83) |
| *Macaca mulatta* | Rhesus macaque | 10.5 (84) | 12 (82) | 3.5 (83) |
| *Papio anubis* | Baboon | 11 (85) | 11 (66) | 5.4<sup>+</sup> (86) |
| *Cercopithecus aethiops* | Green Monkey | 10.2 (87) | 11<sup>+</sup> (88) | 5 (83) |
| *Saimiri sciureus* | Squirrel Monkey | 10.2 (87) | 9* (89) | 3 (90) |
| *Callithrix jacchus* | Marmoset | 10 (91) | 6 (20) | 0.9 (92) |

Note: n/a = not available. \* only female generation was available. <sup>+</sup> inferred from a closely related species.
<sup>a</sup> source: (22)
<sup>b</sup> main source: (90)

**Table S3:** Proportion of substitution in each context across lineages.

| Species | A/T transitions | A/T transversions | G/C transitions | G/C transversions | CpG transitions | CpG transversions | non-CpG G/C transitions | non-CpG G/C transversions |
| --- | --- | --- | --- | --- | --- | --- | --- | --- |
| hg19 | 0.344 | 0.152 | 0.340 | 0.164 | 0.118 | 0.014 | 0.221 | 0.150 |
| panTro4 | 0.333 | 0.163 | 0.339 | 0.164 | 0.116 | 0.012 | 0.222 | 0.151 |
| hg19-panTro4 | 0.334 | 0.150 | 0.357 | 0.158 | 0.119 | 0.013 | 0.237 | 0.145 |
| ponAbe2 | 0.319 | 0.145 | 0.373 | 0.162 | 0.112 | 0.012 | 0.259 | 0.150 |
| hg19-ponAbe2 | 0.303 | 0.160 | 0.380 | 0.157 | 0.118 | 0.014 | 0.261 | 0.142 |
| rheMac3 | 0.289 | 0.170 | 0.377 | 0.164 | 0.124 | 0.013 | 0.252 | 0.150 |
| macFas5 | 0.302 | 0.162 | 0.387 | 0.149 | 0.136 | 0.014 | 0.251 | 0.135 |
| rheMac3-macFas5 | 0.315 | 0.158 | 0.383 | 0.144 | 0.123 | 0.014 | 0.259 | 0.130 |
| papHam1 | 0.301 | 0.156 | 0.399 | 0.144 | 0.121 | 0.013 | 0.277 | 0.131 |
| rheMac3-papHam1 | 0.357 | 0.159 | 0.355 | 0.129 | 0.111 | 0.012 | 0.244 | 0.117 |
| chlSab1 | 0.306 | 0.167 | 0.382 | 0.145 | 0.112 | 0.013 | 0.268 | 0.131 |
| rheMac3-chlSab1 | 0.320 | 0.176 | 0.357 | 0.148 | 0.091 | 0.012 | 0.264 | 0.135 |
| hg19-rheMac3 | 0.311 | 0.178 | 0.358 | 0.152 | 0.106 | 0.016 | 0.251 | 0.136 |
| calJac3 | 0.321 | 0.185 | 0.345 | 0.149 | 0.096 | 0.014 | 0.246 | 0.134 |
| saiBol1 | 0.332 | 0.182 | 0.340 | 0.146 | 0.087 | 0.013 | 0.250 | 0.132 |
| calJac3-saiBol1 | 0.291 | 0.193 | 0.365 | 0.152 | 0.082 | 0.014 | 0.280 | 0.137 |

**Table S4:**
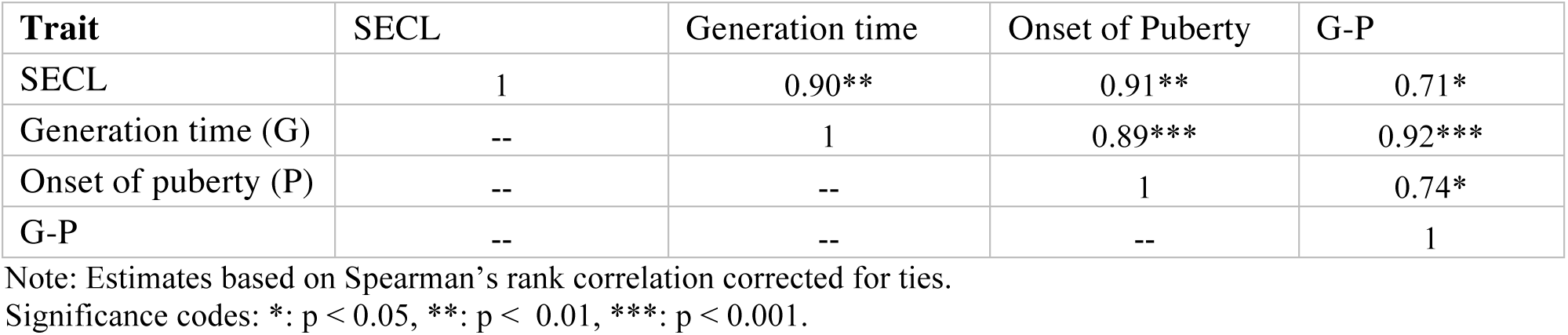
Correlation in life history traits across primates.

| Trait | SECL | Generation time | Onset of Puberty | G-P |
| --- | --- | --- | --- | --- |
| SECL | 1 | 0.90** | 0.91** | 0.71* |
| Generation time (G) | -- | 1 | 0.89*** | 0.92*** |
| Onset of puberty (P) | -- | -- | 1 | 0.74* |
| G-P | -- | -- | -- | 1 |

Note: Estimates based on Spearman's rank correlation corrected for ties.
Significance codes: \*: $p < 0.05$ , \*\*: $p < 0.01$ , \*\*\*: $p < 0.001$ .

